# Single-Molecule Orientation Imaging Reveals the Nano-Architecture of Amyloid Fibrils Undergoing Growth and Decay

**DOI:** 10.1101/2024.03.24.586510

**Authors:** Brian Sun, Tianben Ding, Weiyan Zhou, Tara S. Porter, Matthew D. Lew

## Abstract

Amyloid-beta (Aβ42) aggregates are characteristic signatures of Alzheimer’s disease, but probing how their nanoscale architectures influence their growth and decay remains challenging using current technologies. Here, we apply time-lapse single-molecule orientation-localization microscopy (SMOLM) to measure the orientations and rotational “wobble” of Nile blue (NB) molecules transiently binding to Aβ42 fibrils. We quantify correlations between fibril architectures, measured by SMOLM, and their growth and decay visualized by single-molecule localization microscopy (SMLM). We discover that stable Aβ42 fibrils tend to be well-ordered, signified by well-aligned NB orientations and small wobble. SMOLM also shows that increasing order and disorder are signatures of growing and decaying Aβ42 fibrils, respectively. We also observe SMLM-invisible fibril remodeling, including steady growth and decay patterns that conserve β-sheet organization. SMOLM reveals that increased heterogeneity in fibril architectures is correlated with more dynamic remodeling and that large-scale fibril remodeling tends to originate from local regions that exhibit strong heterogeneity.

## MAIN TEXT

Amyloid diseases such as Alzheimer’s Disease (AD) are incurable neurodegenerative ailments afflicting many older individuals. Amyloid diseases are at least partially caused by protein misfolding, which induces formation of highly insoluble β-sheet fibrils and oligomers that are associated with cytotoxicity.^1–4^ Amyloid-beta peptides such as Aβ42 are thought to comprise most such plaques in AD patients;^5^ consequently, quantifying the diversity of nanometer-size Aβ42 aggregate dynamics and structures is of critical interest to biomedical researchers.^6^

High-speed atomic force microscopy,^7–9^ fluorescence microscopy and spectroscopy^10–12^, and single-particle tracking^13^ have all been used to interrogate how the nanoscale morphologies of amyloid structures change over time. In addition, super-resolved single-molecule localization microscopy (SMLM) has become popular for amyloid imaging. Both binding activated localization microscopy (BALM)^14^ and transient amyloid binding (TAB) imaging^15^ utilize amyloidophilic dyes, such as p-TFAA, Thioflavin T (ThT), Thioflavin X^16^, Nile red, SYPRO orange, and LDS772^17^, whose fluorescence quantum yields and/or emission spectra change when the dyes are in proximity of amyloid fibers. These forms of SMLM can image amyloid aggregates for hours to days without immunostaining or covalently modifying the peptides themselves.^15,18^ While these techniques all achieve high-resolution imaging of aggregate morphology at various temporal resolutions, one key weakness is their inability to resolve the organization or architecture of the peptide assemblies themselves. Such data would, for example, resolve how peptide organization within amyloid aggregates impact their growth, decay, and changes in shape.

To address these weaknesses, one may augment SMLM with spectroscopic capabilities^19–22^ or measure fluorophores’ positions and orientations via single-molecule orientation localization microscopy (SMOLM).^23–26^ SMOLM has been successfully applied to study actin filaments,^27–29^ molecular motors,^30–32^ DNA conformations,^33,34^ and membrane composition and fluidity.^35–37^ By combining the TAB labeling technique with SMOLM, it is possible to visualize how the nanoscale organization of amyloid assemblies affects the orientations of dye molecules bound to their surfaces. In this way, SMOLM has resolved nanoscale structural heterogeneities in amyloid fibers^38,39,40^ and distinct dye binding configurations dependent on amyloid fibril orientation.^17^ Despite these developments, the precise links between the dye orientations sensed by SMOLM, which we call the amyloid fiber’s orientation signature, and their structural dynamics remains unknown.

In this work, we seek to understand how the order and rigidity of fibril β-sheet assemblies, as revealed by the orientations of amyloidophilic dyes, correlate with fiber remodeling over time. First, we introduce two SMLM-based metrics, chi-square distance *χ*^2^ and scaled localization difference *η*, to track changes in Aβ42 fiber shape and the density of hydrophobic binding sites on fiber surfaces over time. We then correlate TAB SMOLM measurements with *χ*^2^ and *η* to quantify the relationships between orientation signatures and Aβ42 growth and decay, respectively. Finally, we utilize TAB SMOLM to measure sub-structural changes in Aβ42 fibers that are invisible to SMLM, thereby demonstrating how SMOLM provides insight into the structural organization and heterogeneity of Aβ42 aggregates that impact their growth and decay.

To begin, we prepared fibrils of amyloid beta as described previously^40^ and adsorbed them to ozone-cleaned cell culture chambers (see SI for details). We utilized a time-lapse TAB^15^ imaging protocol to visualize the remodeling of each amyloid fiber with nanoscale resolution. Specifically, we added the solvatochromic and fluorogenic dye Nile blue (NB)^41–43^ to the imaging chambers after amyloid adsorption (Figure 1a), and “flashes” of NB fluorescence commensurate with binding to the fibril surfaces were captured by an x- and y-polarized microscope^40^ (Figure S1). Typically, 10,000 images of NB blinking were recorded for each SMOLM reconstruction with a 20 ms exposure time under 637-nm illumination. A bespoke regularized maximum-likelihood estimator (RoSE-O) was then used to estimate the 2D position (x,y), in-plane orientation *ϕ*, and rotational "wobble” Ω of the NB molecule (Figure 1a) associated with each flash.^44,45^ We quantify wobble via the solid angle Ω of the hard-edged cone within which a molecule diffuses during the camera’s integration time.

**Figure 1:**
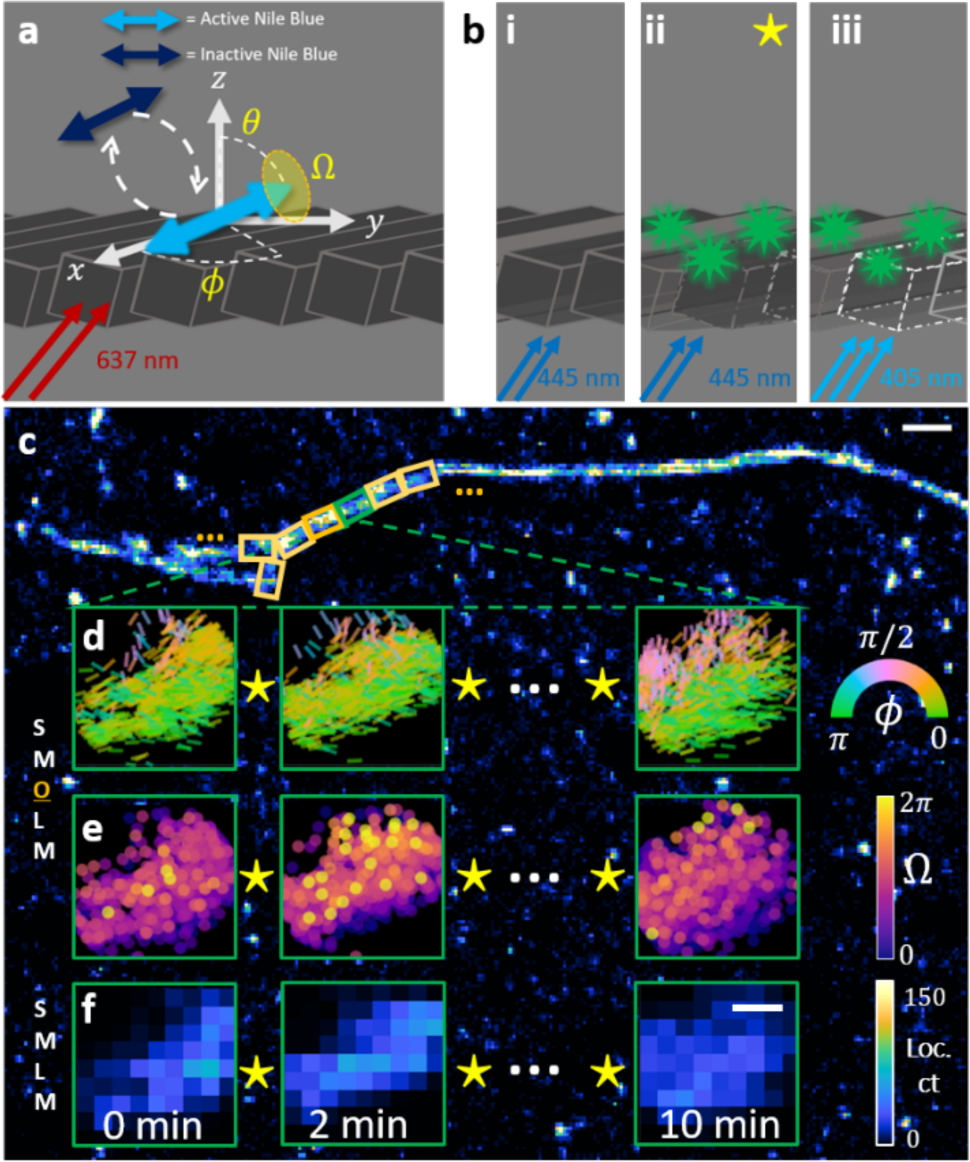
Time-lapse single-molecule orientation-localization microscopy (SMOLM) of Aβ42 remodeling. **(a)** Schematic of imaging buffer showing transient amyloid binding (TAB) of NB (double-headed arrows) to an amyloid beta-42 (Aβ42) fiber (cross-beta sheets shown as gray rectangular prisms). Excitation by 637-nm light facilitates “flashes” of Nile blue (NB) fluorescence during binding to Aβ42. From the x- and y-polarized fluorescence images, SMOLM measures NB’s polar *θ* and azimuthal *ϕ* orientation and rotational wobble within a cone of solid angle Ω. **(b)** Schematic of remodeling buffer, where the concentration of thioflavin T (ThT, green stars) and wavelength and intensity of the excitation light are varied (i-iii) to induce a range of Aβ42 remodeling behaviors (Table S1). **(c)** Single-molecule localization microscopy (SMLM) image of an Aβ42 fiber before remodeling; yellow boxes denote 200 nm-long regions of interest (ROIs). Each Aβ42 fiber undergoes time-lapse imaging over several minutes using the imaging configuration in a, with remodeling (⋆) induced between each imaging step using a buffer condition in b. **(d, e)** SMOLM measurements of NB **(d)** orientation *ϕ* and **(e)** wobble Ω within a selected ROI (green box in c). NB orientations *ϕ* are represented as color-coded line segments aligned parallel to the measured orientation. **(f)** NB localization counts via SMLM. Colorbars: (d) NB orientation *ϕ* (rad), (e) NB wobble Ω (sr), (f) NB localizations per 20 nm × 20 nm bin. Scalebars: (c) 300 nm, (d-f) 60 nm.

To facilitate Aβ42 remodeling, we replace the NB imaging buffer with PBS containing Thioflavin T (ThT, Figure 1b). While ThT, ThX, and other relatives are useful TAB imaging agents themselves,^16^ ThT is also known to perturb Aβ42 fibers via the generation of reactive oxygen species when irradiated.^46^ We used 405-nm and 445-nm lasers at various intensities to induce ThT-induced remodeling of Aβ42 (Table S1). A full experiment involves introducing the NB imaging buffer, collecting SMOLM data, replacing the imaging buffer with the remodeling buffer, incubating the sample, and then repeating this process to quantify changes to the fiber architecture over time.

Using NB, TAB SMLM reveals Aβ42 morphologies similar to those seen when using ThT^15^ and Nile red^40^ i.e., extended fibrils with occasional branching (Figure 1c). To quantify these characteristics, we spatially partition each fibril into non-overlapping regions of interest (ROIs) approximately 200 nm in length (Figure 1c inset). Within one example ROI, we observe remarkable changes in the orientations *ϕ* (represented as line segments centered at each NB location and aligned along each NB orientation) and wobble Ω of NB. SMLM and SMOLM can both resolve the morphology of the segment with nanoscale resolution. However, only SMOLM can sense a diversity of NB orientations and wobble even in areas that appear mostly stable in SMLM (Figure 1d,e), suggesting diversity in the structures’ underlying assemblies.

Since TAB SMOLM characterizes individual NB orientations as they bind to amyloid fibrils, we are led to consider a new question: how correlated are NB orientation signatures with structural remodeling of Aβ42 fibers? We begin by defining metrics that robustly quantify SMLM-observed remodeling. Since NB only emits fluorescence in the vicinity of amyloid fibrils, changes in localization density are commensurate with changes in fibril structure. We therefore use chi-square distance *χ*^2^, a commonly used image similarity metric^47^, to compute the change in fiber structure between two SMLM images, where

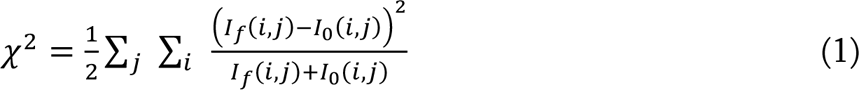

and *I*_0_(*i*, *j*) and *I_f_*(*i*, *j*) represent the normalized NB localization densities of pixel (*i*, *j*) at an initial and final time point, respectively, and the sum is taken over an ROI. We also wish to quantify the total growth and decay of Aβ42 fibers, i.e., whether the number of hydrophobic sites amenable to NB binding is increasing or decreasing. To do so, we define the localization difference metric *η* as

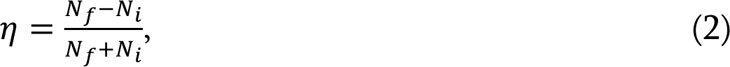

where *N_i_* and *N_f_* represent the initial and final number of localizations, respectively, within an ROI.

We then calculate *χ*^2^ and *η* for all Aβ42 ROIs (see Table S1 for ROI details). As expected of similarity metrics in general, we observe that *χ*^2^ scales monotonically with the extent of remodeling. Smaller *χ*^2^ values correspond to minor shape changes (Figure 2(i-iii)). Regions with *χ*^2^ exceeding ∼0.25 exhibit more significant shape changes (Figure 2iv). Very dramatic morphological changes, such as those seen in unstable or newly formed segments (Figure S2a inset), correspond to *χ*^2^ values exceeding ∼0.5. In contrast, smaller absolute |*η*| values (less than ∼ 0.2) correspond to mostly stable Aβ42 ROIs (Figure 2i,ii). Moderate localization differences |*η*| are associated with a middling degree of growth or decay, while |*η*| values exceeding ∼0.5 are associated with Aβ42 ROIs that grow or decay drastically (Figure 2iii,iv; S2a insets). Thus, both metrics are suitable for quantifying Aβ42 ROI remodeling using SMLM images, and we classify the degree of change (Figure 2c) and amount of growth and decay (Figure 2d) using the aforementioned *χ*^2^ and *η* thresholds.

**Figure 2:**
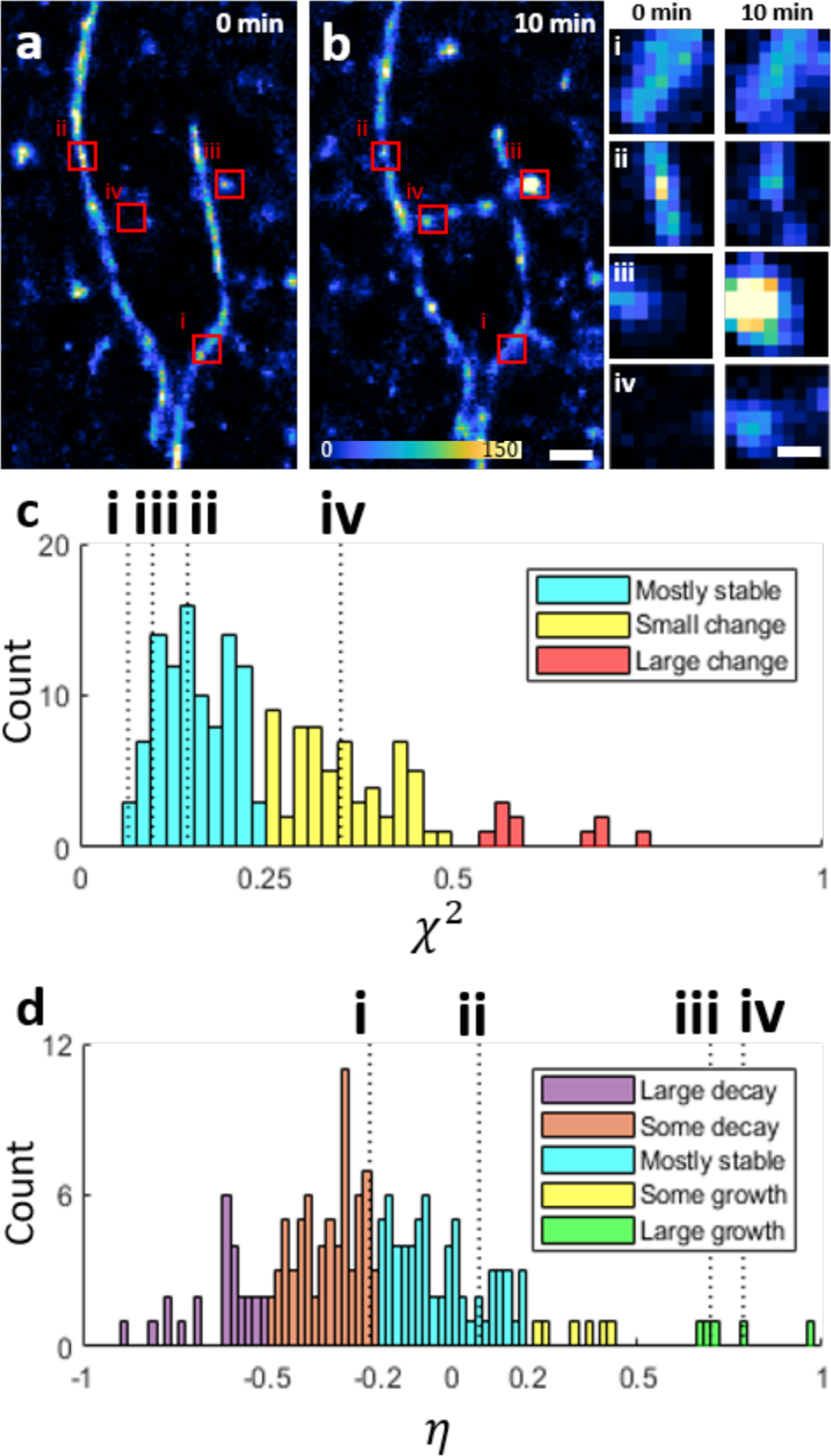
SMLM quantifies amyloid fiber stability and remodeling. **(a,b)** Aβ42 fiber (a) before and (b) after 10 minutes of ThT-induced remodeling. (i-iv) Examples of (i) stable, (ii) decaying, (iii) rapidly growing, and (iv) newly formed segments, indicated by the red boxes. **(c)** Distribution of observed changes in fiber shape (χ^2^), where thresholds χ^2^ = 0.25 and χ^2^ = 0.5 separate the categories of mostly stable, small change, and large change**. (d)** Distribution of observed fiber growth and decay (*η*), where the thresholds |*η*| = 0.2 and |*η*| = 0.5 separate the categories of mostly stable, some growth/decay, and large growth/decay. The specific values of *χ*^2^and *η* for ROIs (i) through (iv) are indicated using dashed lines. Colorbar: NB localizations per 20 nm × 20 nm bin. Scalebars: (a) 300 nm, (i-iv) 80 nm.

We also analyzed the orientation signatures of individual NB fluorophores interacting with amyloid fibrils via SMOLM. We noticed that seemingly stable Aβ42 ROIs appear to have well-aligned orientations (Figure 3a), as indicated by a small standard deviation *σ_ϕ_* in *ϕ* and a small degree of average wobble 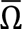 taken across all NB localizations within the ROI (Figure S3a). Both the structures and the measurements remain stable over time. Rapidly growing Aβ42 ROIs, such as the segment pictured in Figure 3b, are associated with decreasing *σ_ϕ_* (Figure S3b), showing that NB orientations become more aligned as the structures and their underlying assemblies become more organized. Rapidly decaying Aβ42 ROIs, such as in Figure 3c, are associated with increasing *σ_ϕ_* and 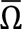 (Figure S3c), thereby illustrating that NB binds with more varied orientations and less rigidity as the structures degrade.

**Figure 3:**
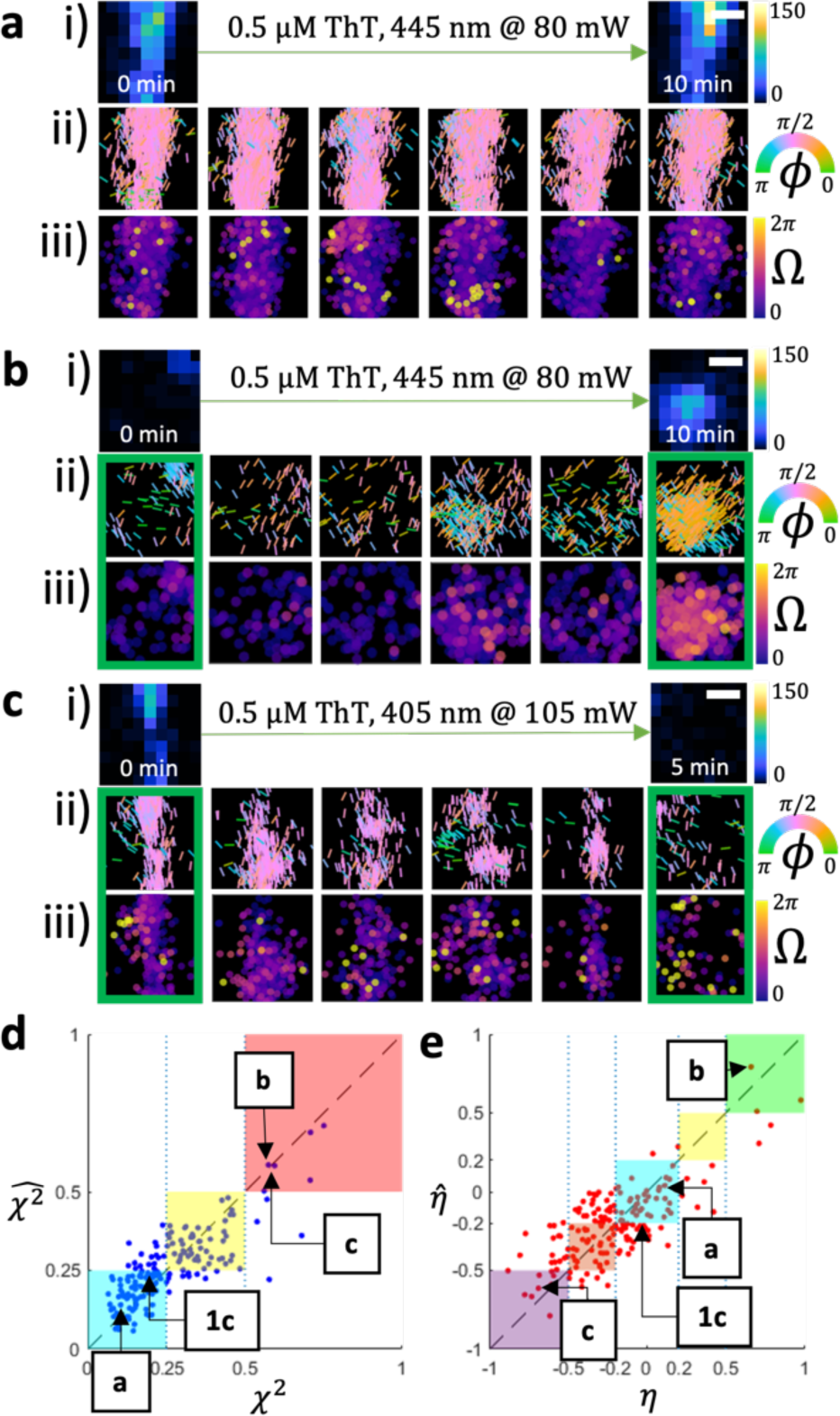
SMOLM quantification of fibril remodeling, growth, and decay. **(a)** Stable Aβ42 segment measured over 10 minutes. **(i)** SMLM and SMOLM images of **(ii)** orientation *ϕ* and **(iii)** wobble Ω. The orientation statistics of the segment remain stable over time (see Fig. S3a for details). **(b)** Same as (a) but for a growing Aβ42 oligomer imaged over 10 minutes. Note the increased organization of the oligomer in the final timestamp compared to all preceding timestamps, as quantified by *ϕ* (boxed in green, see Fig. S3b for details). **(c)** Same as (a) but for a rapidly decaying Aβ42 segment imaged over the course of 5 minutes. Note the large increase in 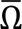 and the increased disorder in *ϕ* between the initial and final timestamps (boxed in green, see Fig. S3c for details). **(d)** Correlation between change in fibril morphology 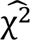 predicted by our regression model from SMOLM data and the change χ^2^ observed by SMLM. The 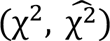 pairs for the segments in a, b, c, and Fig. 1c are indicated. The classification agreement (number of data points within the shaded squares) is 81.9%. **(e)** Correlation between segment growth/decay 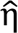 predicted by our regression model from SMOLM data and the growth/decay η observed by SMLM. The 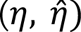 pairs for the segments in a, b, c, and Fig. 1c are indicated. The classification agreement (number of data points within the shaded squares) is 48.5%. Colorbars: (i) NB localizations per 20 nm × 20 nm bin, (ii) NB orientation *ϕ* (rad), (iii) NB wobble Ω (sr). Scalebars: 60 nm.

Given these relationships, we explored whether relationships between NB orientation signatures and fibril architectural dynamics could be modeled using simple polynomial regressions. We computed a series of SMOLM statistics that quantify temporal changes in both average orientation 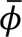 and wobble 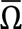 as well as changes in standard deviations *σ_ϕ_* and *σ*_Ω_ within each ROI (Table S2). We then constructed two third-order polynomial regression models to correlate these SMOLM metrics with the SMLM-based measures of fiber remodeling, *χ*^2^ and *η*, retaining only the significantly correlated terms (p < 0.05) in the final models (Supporting Information Section 7, Tables S3 and S4).

We observed a remarkable correlation between SMOLM statistics and the fibril shape changes quantified by *χ*^2^. Aβ42 ROIs whose shapes are mostly stable or experience some changes, such as the segments in Figures 3a, S4a, and S4b, are assigned categories by the SMOLM regression model that largely match those measured by SMLM (83.2% and 82.3% agreement for stable and small changes, respectively, Fig. 3d cyan and yellow). These results indicate that relatively stable Aβ42 regions share similarly stable *ϕ* and Ω signatures, i.e., measured values of 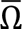, 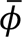, *σ_ϕ_* and *σ_Ω_* that are relatively stable over time. Furthermore, we measured smaller variations in orientation *σ_ϕ_* and wobble *σ_Ω_* for these segments compared to their more dynamic counterparts (Figure S4), suggesting that stable structures tend to exhibit homogeneous binding rigidity for NB and uniform cross-β groove orientations.

However, Aβ42 segments that change drastically, such as those in Figs. 3b and 3c, exhibit less concordance between the SMOLM regression model and SMLM observations (60% agreement, Figure 3d). We attribute this moderate agreement to the diversity of mechanisms that drive large shape changes in Aβ42 fibrils, such as drastic growth (Fig. 3b), drastic decay (Figure 3c), or dramatic reorganizations not accompanied by significant growth or decay (Figure S6). Because growing, stable, and decaying regions are correlated with many possible structural features within amyloid fibrils, we naturally expect a simple polynomial regression to only be partially successful at correlating diverse SMOLM statistical trends with morphological dynamics observed by SMLM.

Correlations between SMOLM statistics and SMLM measurements of localization differences *η* (Figure 3e) also exist, but the relationship is weaker. About half of Aβ42 ROIs exhibiting mostly stable or slight decreases in NB binding activity are correctly identified (59.09% and 46%, Figure 3e cyan and red, respectively). We discover that unchanging (Figure S4a), slightly decaying (Figure S4b), or moderately growing (Figure S4c) Aβ42 structures are all associated with small changes in average orientation 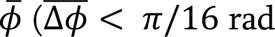, where 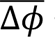 signifies the average magnitude of change in mean orientation between subsequent measurements), small variations in orientation *σ_ϕ_* ( Δ*σ_ϕ_*_,net_ ∈ [−0.1, 0.1] sr and 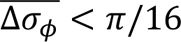 rad, where 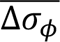 signifies the average magnitude of change in orientation spread between subsequent measurements), small average wobble 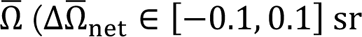 and 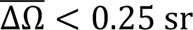, where 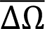 signifies the average magnitude of change in mean wobble between subsequent measurements), and small variations in wobble 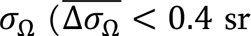, where 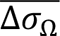 signifies the average magnitude of change in wobble spread between subsequent measurements). As a result, the extent of growth in all moderately growing Aβ42 regions is underestimated (yellow region in Figure 3e). These findings are consistent with our *χ*^2^ correlation analysis; more stable structures tend to exhibit homogeneous binding rigidity and well-aligned β-sheet orientations. Aβ42 structures undergoing significant net growth share common *ϕ* and Ω signatures that also happen to be distinct from other overall growth and decay patterns; most regions with *η* ≥ 0.5 are categorized by SMOLM regression as having drastically grown (60% agreement). However, when estimating large decay, the regression model tends to underestimate the extent (36% concordance, 64% underestimated decay, Figure 3e purple). Again, we attribute these mismatches to the diversity of structural architectures associated with fibril deterioration; for example, segments can decay drastically yet largely maintain their underlying structure (Figure S5a). This pathway yields smaller changes in 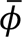 and smaller increases in *σ_ϕ_* - signatures that closely resemble markers of gradually decaying regions. Segments can also decay in more haphazard fashions that greatly disrupt the underlying β-sheet organization, as seen in the segments shown in Figures 3c and S5b. These dynamics are reflected in large net changes in 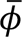 and large net increases in *σ_ϕ_* and 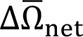, all of which sufficiently distinguish the ROIs as undergoing drastic decay. In totality, we find that simple polynomial regression models trained using amyloidophilic dye orientations from SMOLM correlate moderately with SMLM measurements of fibril remodeling (81.9% agreement) and growth and decay (48.5% agreement).

We also discovered that TAB SMOLM senses nanoscale structural dynamics that are invisible to SMLM, e.g., changes in fiber architecture not associated with large *χ*^2^. To quantify these changes hidden to SMLM, we split Aβ42 ROIs into smaller 50 nm × 50 nm subROIs and quantify time-averaged NB SMOLM statistics in each subROI, e.g., average orientational standard deviation 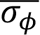 and average wobble 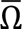. We then compare each subROI to its neighbors, as well as against themselves at different points in time, to measure both spatial and temporal correlations in these signatures.

First, we observe that disorganized structures (Figures 4 and S7-S9, dashed yellow boxes) tend to be more disordered than their more uniform neighbors (Figures 4 and S7-S9, solid green boxes), as reflected in higher 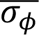 and 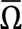 measurements (Figures S10 and S11). These trends are also observed when NB binding is sparse; the edges of the fibril in Figures 4b(i-iii) exhibit higher 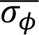 and 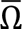 than their more well-defined neighbors in the core of the segment (Figures 4b(iv-vi), see Figures S10b and S11b for precise statistics). This correlation suggests that when NB encounters a high concentration of binding sites oriented in many directions, NB also wobbles to a large degree as it explores them. Moreover, disorganized structures are also significantly more dynamic. For example, the assemblies in Figure 4a(i-iii) show larger 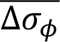 and 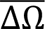 (Figures S10a and S11a) than their more uniform neighboring subROIs (Figure 4a(iv,v)). Similarly, exposed edges of the Aβ42 fibrils (Figures 4b(i-iii) and 4c(iii,iv)) exhibit not only higher 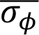 and 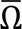 compared to the cores of the fibrils but also larger average fluctuations (Figures S7, S8, S10b, S10c). In general, we observed that NB molecules exhibit larger wobble when binding to disorganized Aβ42 structures. Further, SMOLM reveals that these structures exhibit much larger fluctuations in their architectures than their more uniform counterparts (Figure S10, yellow vs. green subROIs).

**Figure 4:**
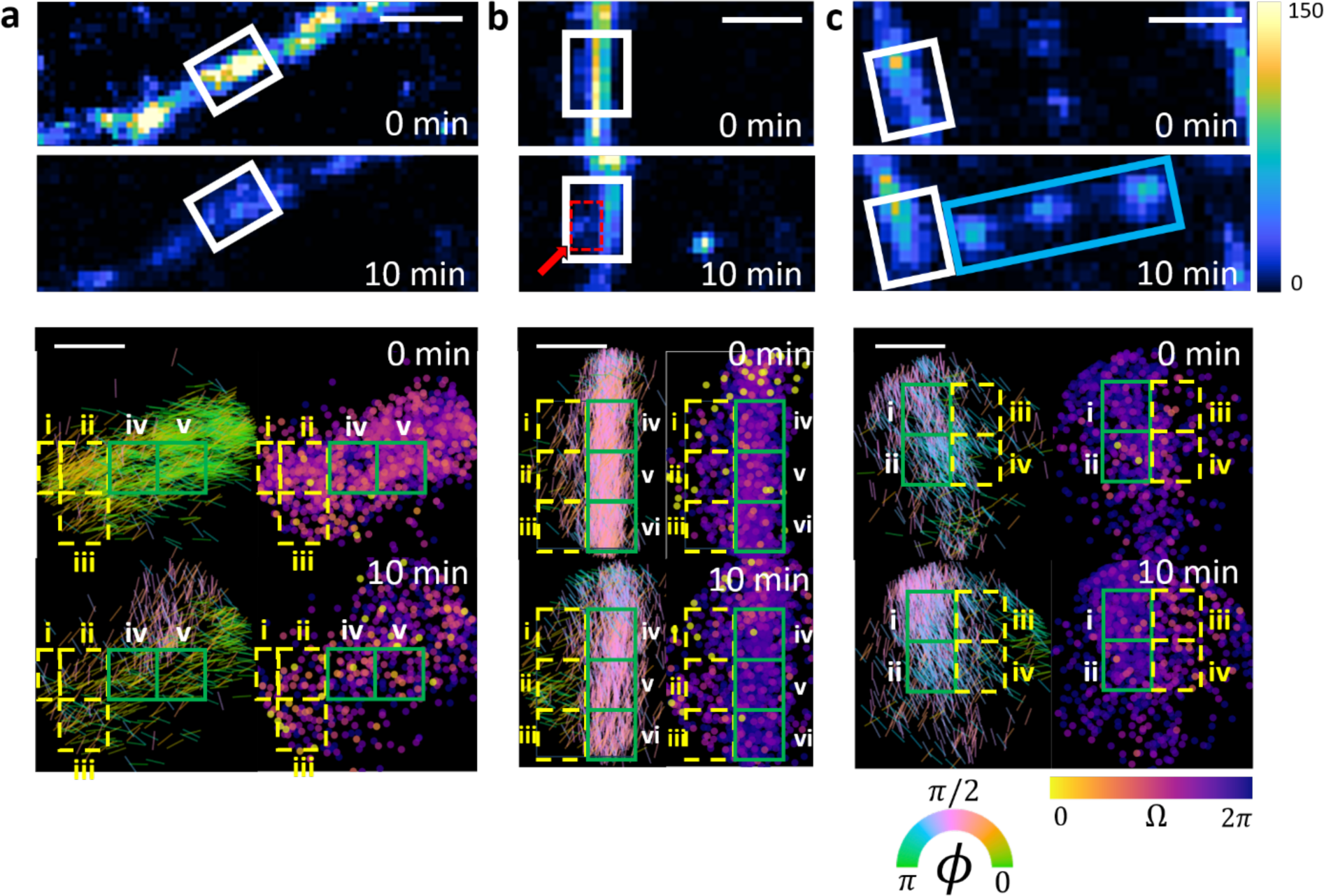
SMOLM reveals architectural remodeling at the nanoscale. **Top:** SMLM images taken at 0 and 10 min. **Bottom:** SMOLM images of the subROI within white box (top). (Left) Orientation *ϕ* of each NB molecule, represented as a line segment aligned parallel to its orientation. (Right) Wobble Ω of each NB molecule. **(a)** Linear segment remodeling over 10 min. SubROIs (i-iii) (dashed yellow boxes) feature more disordered underlying assemblies on average than their neighboring subregions (iv,v) (green boxes). NB shows weaker binding to heterogeneous subROIs (i-iii) compared to neighbors in the core of the fiber (iv,v). Only small changes in segment shape are observed by SMLM (*χ*^2^ = 0.129) in subROIs (i-iii). **(b)** Left edges of a linear segment remodel over 10 min. SubROIs (i-iii) (dashed yellow boxes) have more disordered underlying assemblies than the core of the fibril (iv-vi) (boxed in green). The fringes are also more structurally heterogeneous than their neighbors in the segment core. The growth of a disorganized offshoot in subROI (ii) (red dashed box at 10 min.) is associated with only a small change in overall shape (*χ*^2^ = 0.082). **(c)** A linear segment with a heterogenous right section remodeling over 10 min., culminating in the formation of a rightward “bridgehead” structure (cyan box). The dynamic edges of the fibril in subROIs (iii,iv) (dashed yellow boxes) are more disordered and structurally heterogeneous than the segment core in subROIs (i,ii) (green boxes), with more dynamic changes in underlying assembly orientation and rigidity in the fringes compared to the core. Colorbars: (top, SMLM) NB localizations per 20 nm × 20 nm bin, (bottom left, SMOLM) NB orientation *ϕ* (rad), (bottom right, SMOLM) NB wobble Ω (sr). Scalebars: (top, SMLM) 200 nm, (bottom, SMOLM) 80 nm.

Most strikingly, we observed dramatic seeding of new aggregates by these same disorganized subregions via SMLM, often minutes after SMOLM detects changes in nano-architecture. A disorganized offshoot (yellow arrow and dashed boxes in Figure 4b) grows from disorganized adjacent subregions (Figure 4b(i-iii)), as seen at 2 and 4 min. in Figure S8. More strikingly, an oligomeric “bridge” (cyan box in Figure 4c) grows between two arms of an Aβ42 fibril; this bridge appears to be seeded by a “bridgehead” subregion with a high degree of disorganization (Figure 4c(iii,iv)), as seen at 6 and 8 min. in Figure S9. These data suggest that SMOLM can detect changes in fibril nano-architecture, manifesting as disorganized structures on the ∼50-nm scale, well before SMLM detects dramatic remodeling.

In summary, we utilize a minimally perturbative labeling approach, the transient amyloid binding (TAB) of fluorogenic Nile blue molecules, to quantify the structural architecture and remodeling of amyloid-beta fibrils using SMOLM. We demonstrate two SMLM-based metrics for quantifying fibril remodeling (*χ*^2^) as well as growth and decay (*η*). We then correlate, via polynomial regression, SMOLM statistics of NB orientations to measures of Aβ42 remodeling (Figure 3d,e). We find that ordered and uniformly oriented assemblies correspond to stable Aβ42 structures (Figures 3a and S4), that assemblies increasing in uniformity tend to grow (Figures 3b and S3b), and that increasingly disordered assemblies correspond to decaying structures (Figures 3c and S5). Finally, we use TAB SMOLM to reveal that locally disordered regions can facilitate large-scale fibrillar remodeling (Figures 4 and S7-S11).

We also observe SMLM measurements of growth, decay, and remodeling that do not correlate with SMOLM measurements of NB orientations and their dynamics. TAB SMOLM shows that Aβ42 remodeling processes are nuanced; Aβ42 structures can steadily grow and decay while retaining their underlying β-sheet architecture, with associated NB orientation measurements that consequently resemble stable Aβ42 structures. Aβ42 structures can also dramatically reorganize their underlying β-sheet assemblies without significant growth or decay, commensurate with NB orientation measurements that suggest drastic remodeling (Figure S6). These behaviors are to be expected, as many mechanisms exist for amyloid-beta growth, decay, and remodeling that are influenced by the many possible pathways and associated kinetics of amyloid formation. Amyloid-beta can form oligomeric structures, organize into fibrils like the ones analyzed in this study,^48^ and alternate between myriad other folded, partially folded, and unfolded states.^49^ Remarkably, SMOLM has elucidated one such pathway of oligomeric bulb growth, i.e., the formation of “bridgeheads” in dynamic subregions that seed oligomer formation (Figure 4b,c).

Our study provides structural signatures of amyloid-beta remodeling using orientations of amyloidophilic imaging dyes. We anticipate that recent advances in utilizing fluorogenic dyes for SMOLM will enable improved understanding of amyloid aggregation. Correlated multi-dye studies,^17,42^ where each fluorophore has unique binding behaviors and/or environmental sensitivity, could elucidate new nanoscale insights into amyloid structure when combined with SMOLM. Through careful analysis of the SMOLM data, structural polymorphisms in the β-sheet assemblies may be resolvable.^38^

Beyond our proof-of-principle experiments, different Aβ42 seedings can be utilized to prepare and analyze a selection of pathologically relevant Aβ42 structures, thereby enabling the study of how different preparations affect structural dynamics. More broadly, future studies can also target other amyloid peptides implicated in neurodegeneration, such as Aβ40. Dynamic SMOLM measurements can also be correlated with ultrahigh resolution characterization of amyloid-beta structures using cryogenic electron microscopy and transmission electron microscopy.^50^ The combination of improved imaging precision and fluorogenic probes specific to particular amyloid structural motifs will deepen our understanding of fundamental mechanisms underpinning amyloid-beta dynamics and potentially reveal new targets for anti-amyloid therapeutics.

## Supporting information

Movie S3: Decaying A-beta 42 segment (Fig. 3c)

Movie S2: Growing A-beta 42 segment (Fig. 3b)

Movie S1: Stable A-beta 42 segment (Fig. 3a)

## ASSOCIATED CONTENT

### Supporting Information

The following files are available free of charge.

Details on the preparation of aggregates, imaging procedures, image and data analysis, regression, and additional examples are provided as a supporting information PDF. Animations of amyloid remodeling are provided in Movies S1-S3 (MP4).

## AUTHOR INFORMATION

### Author Contributions

BZ, TD, and MDL conceptualized the study. BS, TD, and MDL designed the methodology. TD and TP prepared samples and collected data. BS, TD, and WZ developed software used in the study. BS, TD, and WZ analyzed the data. BS visualized the data. BS and MDL wrote the manuscript with input from all authors. MDL supervised, managed, and acquired financial support for the project. All authors have given approval to the final version of the manuscript.

## ACKNOWLEDGMENT

The authors thank Tristan Carlson, Yiyang Chen, Nikita Gupta, Ziyi Hu, Yuanxin Qiu, and Tingting Wu for helpful suggestions and discussions.

## ABBREVIATIONS

Aβ42: amyloid-beta 42
NB: Nile Blue
SMOLM: Single-Molecule Orientation Localization Microscopy
SMLM: Single-Molecule Localization Microscopy.

