## Supporting Information for "Single-Molecule Orientation Imaging Reveals the Nano-Architecture of Amyloid Fibrils Undergoing Growth and Decay"

### 1. Preparation of Aβ42 aggregates

We followed the procedure described by Ding et al.[1] for synthesis, purification, storage, and lyophilization of Aβ42 monomers.

To improve sample purity, the lyophilized Aβ42 is dissolved in NaOH, sonicated in a cold water bath and filtered through 0.22 μm and 30 kD centrifugal membrane filters.

Aβ42 fibrils were prepared by incubating monomeric Aβ42 in phosphate-buffered saline (PBS, pH 7.4) at 37°C and sonicated at 200 rpm for 42-50 hours. The aggregated structures are then adsorbed to an ozone-cleaned cell culture chamber (Cellvis, C8-1.5H-N, No. 1.5H, 170 ± 5 μm thickness) one hour after incubation and flushed with PBS to maximize adherence.

### 2. Aβ42 imaging and remodeling buffers

A PBS solution (200 μL) containing 50 nM Nile Blue (Sigma-Aldrich, 370088-1G) was placed into the amyloid-adsorbed chambers for transient amyloid binding (TAB) single-molecule orientation localization microscopy (SMOLM). A PBS solution (200 μL) containing 0.5 μM Thioflavin T (Sigma-Aldrich, T3516-5G) was placed into the amyloid-adsorbed chambers for remodeling. To exchange the buffers, the old buffer was carefully extracted from the chamber, leaving some solution remaining to keep the fibrils hydrated; the chamber was then washed once with clean PBS (200 μL). Finally, new buffer was added to the chamber slowly to minimize disturbance of the fibrils.

### 3. Time-lapse SMOLM protocol

Aβ42 remodeling is induced via photoexcitation of ThT by illuminating it with 405 nm (OBIS 405) or 445 nm (Cobolt 0445-06-01-0100-130) lasers. This procedure is performed over a time interval ranging from 1-4 minutes.

The TAB SMOLM imaging procedure is very similar to the procedure described in Ding et al.[1] (Figure S1). NB is excited using an inclined 637 nm laser (OBIS 637). The laser is aligned at a ~30° tilt from normal to reduce background fluorescence. Fluorescence from blinking NB molecules is collected by a 100× 1.4 NA oil-immersion objective lens (Olympus, UPlan-SApo 100×). Fluorescence is filtered by a dichroic beamsplitter (Semrock, Di03-R637) and a bandpass filter (Semrock, FF01-676/37) followed by separation into two orthogonally-polarized detection channels by a polarizing beamsplitter (Meadowlark Optics, BB-100-VIS).

Both channels were captured by a scientific CMOS camera (Hamamatsu, C11440-22CU) with a pixel size of $58.5\times58.5$ nm$^2$ in object space and a conversion gain of 0.49 ADU/photon. Image stacks of 10,000 frames with 20 ms exposure were recorded.

A full experiment involves introducing the TAB-NB imaging buffer, collecting SMOLM data, replacing the imaging buffer with the remodeling buffer, incubating the sample, and then repeating this process to visualize fiber remodeling over several time points.

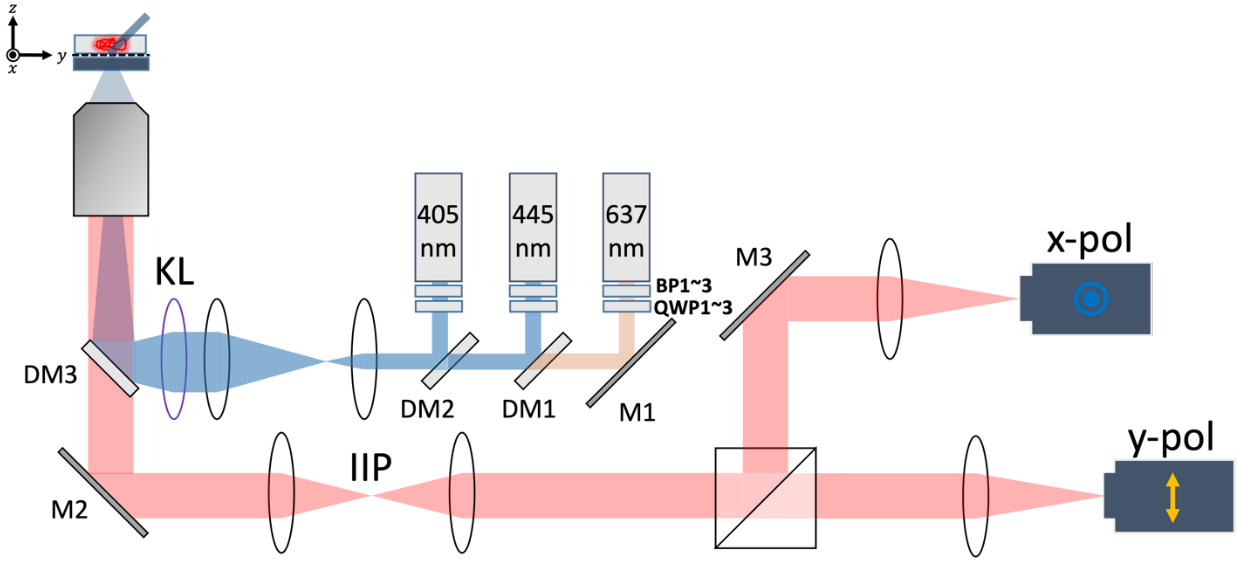

**Figure S1: xy-Polarized Imaging System Schematic.** Circularly polarized 405 nm and 445 nm lasers were used for the excitation of Thioflavin T (ThT), and a circularly polarized 637 nm laser was used for the excitation of Nile Blue (NB). The excitation beam is coupled into a 100X 1.4NA oil objective after expansion via two lenses and a Kohler Lens (KL) and used for inclined sample illumination. Fluorescence is also collected by the objective and filtered through a dichroic mirror (DM3) and bandpass filter before being separated into x- and y-polarized light via a polarized beamsplitter (PBS). Each polarization channel is imaged onto separate regions of an sCMOS camera (two separate cameras are shown for convenience). IIP, intermediate focal plane. DM1-3, dichroic mirrors. M1-3, mirror. BP1-3, bandpass filters. QWP1-3, quarter wave plates.

### 4. Image registration

We follow the image registration procedure described by Ding et al.,[1] which is briefly described as follows. Raw images captured from the xyPol imaging system are offset corrected, and the background photons per pixel are estimated to obtain spatial profiles of the background fluorescence. To properly align the orthogonally polarized fluorescence images simultaneously captured by the CMOS camera, we collect images of spin-coated fluorescent beads, i.e., isotropic point sources. The average localized position of these beads is then obtained using the ThunderSTORM plugin in ImageJ, and coefficients of a 2D polynomial transformation between the points is generated using *fitgeotrans()* in MATLAB. To further remove sample-dependent aberrations, the 2D transformation is refined using bright NB localizations. Only the highest-precision control points (<10 nm localization precision) were used as control points.

### 5. Localization and orientation estimation of NB single molecules

The spatial positions and orientations of fluorescent NB molecules are estimated using a bespoke regularized maximum-likelihood estimator (RoSE-O), as described in Ding et al.[1] The object space is represented as rectangular lattice of grid points with spacing equal to the camera pixel size (58.5 nm), with each molecule parametrized by brightness ($s$), orientational second moments ($\widetilde{m}$), and a positional offset ($\Delta x, \Delta y$) from the grid. RoSE-O then detects all molecules within a raw image and estimates these parameters for each molecule. The orientational second moments are then projected to first moment space (azimuthal orientation $\phi \in [-\pi, \pi)$, vertical orientation $\theta \in \left[0, \frac{\pi}{2}\right]$, and rotational diffusion / "wobble" area $\Omega \in [0, 2\pi]$) by solving a weighted least-squares minimization problem (see Ref. [1] for details).

### 6. Aβ42 ROI segmentation algorithm

SMLM images of fibrils are generated by binning all NB localizations into 20 nm $\times$ 20 nm pixels. We then measure the backbone morphology of each Aβ42 fibril using the Ridge Detection plugin for ImageJ, and all NB localizations exceeding a specific spatial distance from the backbone are excluded as "off-fibril" localizations. Following this filtering process, Aβ42 fibrils are partitioned into spatially isolated regions of interest (ROIs) that are 200 nm in length, and NB localizations are assigned to their nearest ROIs. Doing so enables comparisons of SMOLM signatures within each ROI to both other ROIs and (if desired) the SMOLM signatures of the entire fibril.

**Table S1: Experimental conditions, localization count, and photon number information for imaged Aβ42 fibers and associated segments (ROIs).** Note that an OBIS 637 nm at 60 mW power is used to excite Nile Blue for all fibers. See Figure S2 for SMLM images of all fibers mentioned below.

| Aβ42 fiber (figure #s) | # of Segments (ROIs) | Time lapse images x interval (min) | Remodeling laser (power @ wavelength) | ThT concentration (μM) | Localizations/ segment (avg ± std) | Photons/ localization (avg ± std) |
|---|---|---|---|---|---|---|
| **Horizontal (1c, 4a, S4b, and S5a)** | 39 | 5 x 2 | 100 mW @ 405 nm | 0.5 | 667 ±376 | 3186 ±1766 |
| **Y-shaped (2a, 2b, 3a, 3b, 4b, 4c, S4a, and S4c)** | 40 | 5 x 2 | 80 mW @ 445 nm | 0.5 | 1205 ±608 | 1931 ±1206 |
| **L-shaped (3c and S5b)** | 45 | 5 x 1 | 100 mW @ 445 nm | 0.5 | 441 ±246 | 1658 ±1015 |
| **Curved (S6)** | 47 | 5 x 4 | N/A | 0 | 695 ±387 | 1702 ±1085 |

### 7. Regression models for correlating SMOLM statistics to SMLM measures of fiber remodeling

The eleven SMOLM statistics utilized in the regression model are listed in Table S1 below. Each SMOLM statistic quantifies a different aspect of nanoscale remodeling through either the azimuthal orientation ($\phi$) or dipole rotational diffusion / "wobble" ($\Omega$) of Nile blue, collected over an ROI. Examples of how these statistics change over time for select ROIs are shown in Figs. S3-S6 and S10.

**Table S2: SMOLM statistics and definitions.** The delta notation, e.g., $\Delta\phi$, represents a change in a parameter over time, and the overline notation, e.g., $\bar{\phi}$, represents an average over molecules within an ROI. The subscript "i" denotes a time index over a series of time-lapse images.

| SMOLM statistic | Definition |
|---|---|
| $x_1 = \overline{\Delta\phi} = \frac{1}{N-1}\sum_{i=1}^{N-1}\lvert\bar{\phi}_{i+1} - \bar{\phi}_i\rvert$ | Measures the average absolute change in mean dipole orientation of an ROI between time-lapse images. **Quantifies the degree of change in the ROI's overall orientation.** |
| $x_2 = \lvert\Delta\bar{\phi}_{\text{net}}\rvert = \lvert\bar{\phi}_{\text{final}} - \bar{\phi}_{\text{initial}}\rvert$ | Measures **how much the ROI's overall orientation changed over the course of the experiment.** |
| $x_3 = \overline{\Delta\sigma_\phi} = \frac{1}{N-1}\sum_{i=1}^{N-1}\lvert\sigma_{\phi,i+1} - \sigma_{\phi,i}\rvert$ | Measures the average absolute change in orientation spread of an ROI between time-lapse images. **Quantifies the dynamicity of the ROI's degree of order.** |
| $x_4 = \Delta\sigma_{\phi,\text{net}} = \sigma_{\phi,\text{final}} - \sigma_{\phi,\text{initial}}$ | Measures **the overall change to the ROI's degree of order.** |
| $x_5 = \overline{\Delta\Omega} = \frac{1}{N-1}\sum_{i=1}^{N-1}\lvert\bar{\Omega}_{i+1} - \bar{\Omega}_i\rvert$ | Measures the average absolute change in mean dipole wobble of an ROI changes between time-lapse images. **Quantifies the dynamicity of the ROI's structural rigidity.** |
| $x_6 = \Delta\bar{\Omega}_{\text{net}} = \bar{\Omega}_{\text{final}} - \bar{\Omega}_{\text{initial}}$ | Measures **the change in the ROI's overall rigidity over the course of the experiment.** |

| SMOLM statistic | Definition |
|---|---|
| $x_7 = \overline{\Delta\sigma_\Omega} = \frac{1}{N-1}\sum_{i=1}^{N-1}\lvert\sigma_{\Omega,i+1} - \sigma_{\Omega,i}\rvert$ | Measures how the standard deviation of NB dipole orientations corresponding to an ROI changes between imaging timestamps, on average. **<u>Indicates how drastically the ROI's rigidity heterogeneity changed.</u>** |
| $x_8 = \Delta\sigma_{\Omega,\text{net}} = \sigma_{\Omega,\text{final}} - \sigma_{\Omega,\text{initial}}$ | Measures **<u>the change in heterogeneity of the ROI's binding rigidity over the course of the experiment.</u>** |
| $x_9 = \text{std}(\sigma_\phi) = \sqrt{\frac{1}{N}\sum_{i=1}^{N}(\sigma_{\phi,i} - \overline{\sigma_\phi})^2}$ | Measures the standard deviation of the standard deviation of dipole orientations measured for all time-lapse images. **<u>Quantifies how steadily ROI becomes more or less organized.</u>** |
| $x_{10} = \text{std}(\sigma_\Omega) = \sqrt{\frac{1}{N}\sum_{i=1}^{N}\left(\sigma_{\Omega,i} - \overline{\sigma_\Omega}\right)^2}$ | Measures the standard deviation of the standard deviation of NB wobble at each imaging timestamp. **<u>Quantifies how steadily the ROI's binding rigidity becomes more or less heterogeneous.</u>** |
| $x_{11} = \left(\frac{1}{N}\sum_{i=1}^{N}\frac{1}{\sqrt{s_i}}\right)^{-1}$ | The square-root of the number of photons detected from a Nile blue molecule, averaged over all localizations within the ROI and over all time-lapse images. Signifies **<u>our confidence in the SMOLM measurement</u>**, which is proportional to the inverse of SMOLM measurement precision. |

We construct a vector $x$ with entries $x_i$, where $i \in \{1, \ldots, 11\}$, equal to the values of the SMOLM statistics given in Table S1. The values of each SMOLM statistic are normalized so that their numerical magnitudes are comparable before fitting the regression model.

In designing multivariate, third-order polynomial regression models to correlate these SMOLM statistics, we also include a measurement precision weighting factor $x_{11}$, related to the square-root of the number of photons $\sqrt{s}$ detected during a localization event. For any given localization, the inverse of this weighting term is proportional to the measurement precision of any SMOLM quantity[2], i.e.,

$$\sigma \propto \frac{1}{\sqrt{s}}.$$

For all localizations in an ROI averaged over all time points, we define the average measurement precision as

$$\bar{\sigma} = \frac{1}{N}\sum_{i=1}^{N}\frac{1}{\sqrt{s_i}} = \overline{\left(\frac{1}{\sqrt{s}}\right)}.$$

Since smaller precisions signify better measurement performance, we use the inverse of the average precision as our weight quantity.

We then designed multivariate, third-order polynomial regression models to correlate these normalized SMOLM statistics $x_i$ measured across all ROIs against SMLM measurements of chi-square distance $\chi^2$ and scaled localization difference $\eta$. Second-order cross-terms $(x_i x_j, i \neq j)$ are also included to assess potential correlations between SMLM-quantified Aβ42 remodeling and any joint relationships that exist between the values of $\phi$ and $\Omega$ signatures. To guard against overfitting, we limit the maximum order of any individual term within the polynomial to the third power, i.e., $x_i^3$.

We used the function stepwiselm() in MATLAB to perform stepwise regression of a polynomial containing SMOLM statistics correlated against chi-square distance $\chi^2$ and scaled localization difference $\eta$, retaining only the significantly correlated terms ($p < 0.05$) in the two resulting polynomial models. The model terms and coefficients are given in Tables S2 and S3.

**Table S3: Polynomial model relating SMOLM statistics $x_1, \dots, x_{11}$ to SMLM $\chi^2$.** See Table S1 for the definition of each statistic. The model is given by $\widehat{\chi^2} = \sum_i \text{coefficient}_\text{i} \cdot \text{statistic}_\text{i} + c$ and has an adjusted $R^2 = 0.588$ and MSE of 0.008.

| Normalized SMOLM statistic | Coefficient |
|---|---|
| $x_1$ | 0.031 |
| $x_2$ | -0.004 |
| $x_3$ | -0.044 |
| $x_4$ | -0.011 |
| $x_5$ | 0.051 |
| $x_6$ | -0.004 |
| $x_7$ | -0.012 |
| $x_8$ | 0.015 |
| $x_9$ | 0.072 |
| $x_{10}$ | -0.023 |
| $x_1x_2$ | 0.015 |
| $x_2x_{10}$ | 0.043 |
| $x_4x_8$ | 0.045 |
| $x_4x_{11}$ | -0.024 |
| $x_5x_6$ | 0.038 |
| $x_6x_7$ | 0.030 |
| $x_7x_8$ | -0.032 |
| $x_9x_{11}$ | -0.021 |
| $x_1^2$ | -0.022 |
| $x_5^2$ | -0.038 |
| $x_6^2$ | 0.024 |
| $x_7^2$ | 0.019 |
| $x_{10}^2$ | -0.051 |
| $x_5^3$ | 0.007 |
| $x_6^3$ | -0.007 |
| $x_{10}^3$ | 0.013 |
| $x_{11}$ | 0.007 |
| $x_{11}^2$ | 0.012 |
| $x_{11}^3$ | -0.017 |
| $c$ (intercept) | 0.292 |

**Table S4: Polynomial model relating SMOLM statistics $x_1, \ldots, x_{11}$ to SMLM $\eta$.** See Table S1 for the definition of each statistic. The model is given by $\hat{\eta} = \sum_i \text{coefficient}_\text{i} \cdot \text{statistic}_\text{i} + c$ and has an adjusted $R^2 = 0.491$ and an MSE of 0.047.

| Normalized SMOLM statistic | Coefficient |
|---|---|
| $x_1$ | -0.089 |
| $x_2$ | -0.021 |
| $x_3$ | -0.148 |
| $x_4$ | -0.104 |
| $x_5$ | 0.070 |
| $x_6$ | -0.025 |
| $x_7$ | -0.096 |
| $x_8$ | -0.098 |
| $x_9$ | 0.178 |
| $x_1 x_5$ | -0.071 |
| $x_1 x_6$ | -0.091 |
| $x_1 x_9$ | 0.090 |
| $x_1 x_{11}$ | 0.143 |
| $x_2 x_4$ | 0.063 |
| $x_2 x_5$ | 0.098 |
| $x_4 x_6$ | -0.091 |
| $x_5 x_8$ | 0.078 |
| $x_1^2$ | 0.042 |
| $x_2^2$ | 0.037 |
| $x_4^2$ | -0.047 |
| $x_5^2$ | 0.015 |
| $x_7^2$ | -0.044 |
| $x_4^3$ | 0.028 |
| $x_5^3$ | -0.022 |
| $x_7^3$ | 0.029 |
| $x_{11}$ | 0.106 |
| $x_{11}^2$ | 0.175 |
| $x_{11}^3$ | -0.095 |
| $c$ (intercept) | -0.282 |

### 8. Additional examples of remodeling heterogeneity

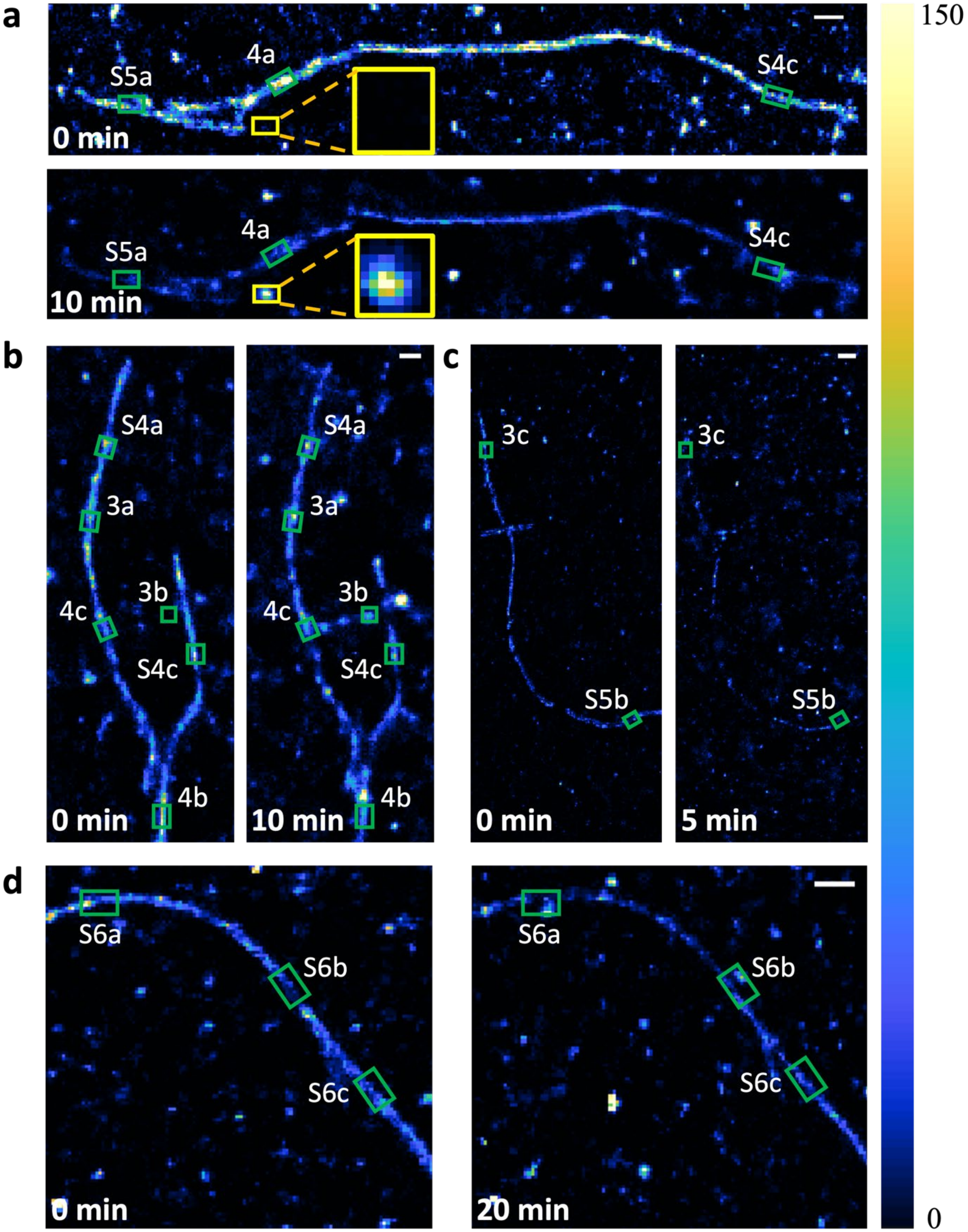

**Figure S2: SMLM images of remodeling Aβ42 fibers. (a)** Fiber from Figure 1c of the main text with segments in Figures 4a, S4b, and S5a indicated in green and a growing oligomer indicated in yellow insets. The oligomer is an example of an Aβ42 segment experiencing abrupt morphological change ($\chi^2 = 0.708$) and growth ($\eta = 0.978$). **(b)** Remodeling Y-shaped fiber with segments in Figures 3a, 3b, 4b, 4c, S4a, and S4c indicated in green. **(c)** Remodeling L-shaped fiber with segments in Figures 3c and S5b indicated in green. **(d)** Remodeling curved fiber with segments in Figure S6 indicated in green. Colorbar: NB localizations per $20\ \mathrm{nm} \times 20\ \mathrm{nm}$ bin. Scalebar: 200 nm.

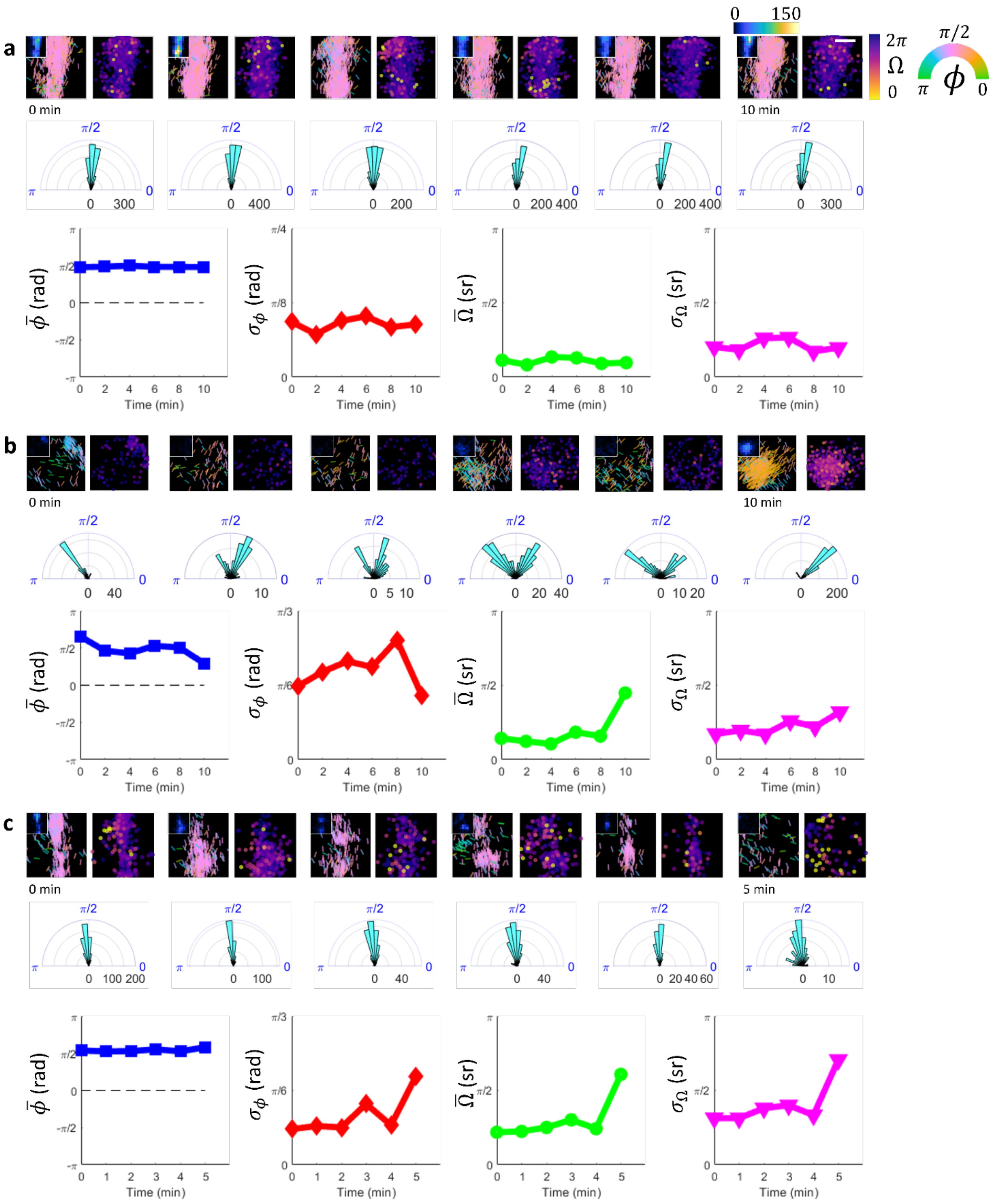

a
b
c
0 min
10 min
5 min
0
150
2π
Ω
0
π/2
π
ϕ
0
Time (min)

**Figure S3: SMOLM analysis of Aβ42 segments from Figure 3**. **(a)** Stable Aβ42 segment associated with stable SMLM and SMOLM statistics imaged over the course of 10 minutes. Note the stable segment morphology ($\chi^2 = 0.071$) and surface hydrophobicity ($\eta = 0.073$). These remodeling changes are associated with small changes in assembly orientation ($|\Delta\phi_{\text{net}}| = 0.009$ rad), uniformity ($\Delta\sigma_{\phi,\text{net}} = -0.014$ rad), wobble ($\Delta\Omega_{\text{net}} = -0.052$ rad), and wobble spread ($\Delta\sigma_{\Omega_{\text{net}}} = -0.016$ rad). As an archetypal example of stability, the extent of remodeling is estimated very well ($\widehat{\chi^2} = 0.143, \hat{\eta} = 0.009$). The trends in average orientation ($\bar{\phi}$), orientation spread ($\sigma_{\phi}$), average wobble ($\bar{\Omega}$), and wobble spread ($\sigma_{\Omega}$) are also shown, highlighting the SMOLM statistics' temporal stability. **(b)** Growing Aβ42 oligomer imaged over the course of 10 minutes. Note the large change in segment morphology ($\chi^2 = 0.574$) and surface hydrophobicity ($\eta = 0.661$) as the oligomer develops. In doing so, the average orientation of the underlying assemblies shifts significantly ($|\Delta\phi_{\text{net}}| = 1.145$ rad), and the assemblies also become more slightly more uniform ($\Delta\sigma_{\phi,\text{net}} = -0.062$ rad). Given the large change in average orientation in combination with the slight increase in assembly organization, the extent of remodeling is estimated well ($\widehat{\chi^2} = 0.585$, $\hat{\eta} = 0.797$). **(c)** Rapidly decaying Aβ42 segment imaged over the course of 5 minutes. Note the rapid loss of segment definition ($\chi^2 = 0.594$) and large decreases in surface hydrophobicity ($\eta = -0.69$), both of which are associated with loss of assembly uniformity ($\Delta\sigma_{\phi,\text{net}} = 0.371$ rad) and large average wobble ($\bar{\Omega} = 1.224$ rad) and increase in wobble spread ($\Delta\sigma_{\Omega,\text{net}} = 1.232$ rad). As an archetypal example of decay, the extent of remodeling is estimated very well ($\widehat{\chi^2} = 0.582, \hat{\eta} = -0.614$).

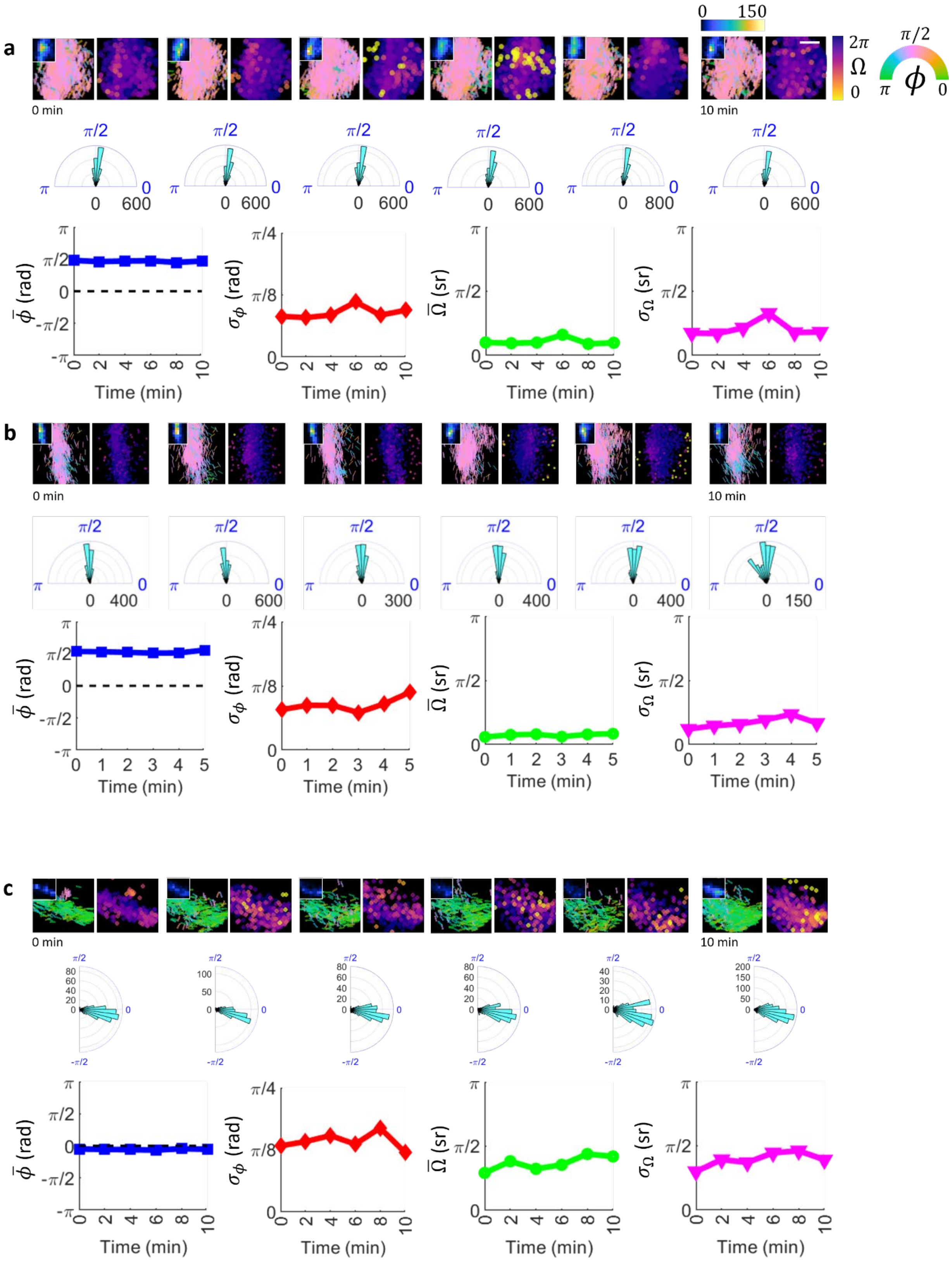
a
b
c
0 min
10 min
0
150
2π
Ω
0
π/2
π
ϕ
0
π/2
π
0
0 600
0 800
0 400
0 300
0 150
π/2
-π/2
80
60
40
20
100
50
200
150
30
10
π
π/2
0
-π/2
-π
π/4
π/8
$\bar{\phi}$ (rad)
$\sigma_\phi$ (rad)
$\bar{\Omega}$ (sr)
$\sigma_\Omega$ (sr)
Time (min)

**Figure S4: SMOLM analysis of relatively stable Aβ42 segments**. **(a)** Stable Aβ42 segment associated with stable SMLM and SMOLM statistics imaged over the course of 10 minutes. Note the stable segment morphology ($\chi^2 = 0.082$) and surface hydrophobicity ($\eta = -0.067$). These remodeling changes are associated with stable, highly ordered β-sheet assemblies as quantified by changes in average NB orientation and spread ($\Delta\sigma_{\phi,\text{net}} = 0.039$ rad and $|\Delta\bar{\phi}_{\text{net}}| = 0.034$ rad). As this segment is an archetypal example of stability, the extent of remodeling is estimated well ($\widehat{\chi^2} = 0.129, \hat{\eta} = -0.15$). The trends in average orientation ($\bar{\phi}$), orientation spread ($\sigma_\phi$), average wobble ($\bar{\Omega}$), and wobble spread ($\sigma_\Omega$) are also shown, highlighting the SMOLM statistics' temporal stability. **(b)** Second stable Aβ42 segment associated with stable SMLM and SMOLM statistics imaged over the course of 10 minutes. As with (a), note the stable segment morphology ($\chi^2 = 0.098$) and segment surface hydrophobicity ($\eta = -0.161$) that are associated with stable, highly ordered β-sheet assemblies as quantified by changes in average NB orientation and spread ($\Delta\sigma_{\phi,\text{net}} = 0.106$ rad and $|\Delta\bar{\phi}_{\text{net}}| = 0.071$ rad), as well as changes in wobble ($\Delta\sigma_{\phi,\text{net}} = +0.149$ sr and $\Delta\bar{\Omega}_{\text{net}} = 0.079$ sr). As another archetypal example of stability, the remodeling extent is also estimated well ($\widehat{\chi^2} = 0.081, \hat{\eta} = -0.133$). **(c)** Moderately growing Aβ42 segment associated with less stable SMLM statistics and stable SMOLM statistics, imaged over the course of 10 minutes. SMLM senses noticeable morphological changes ($\chi^2 = 0.251$) and increasing surface hydrophobicity ($\eta = 0.423$) that are correlated with moderate changes in average NB wobble ($\Delta\bar{\Omega}_{\text{net}} = +0.404$ sr) but highly stable NB average orientation and spread measurements ($|\Delta\bar{\phi}_{\text{net}}| = 0.015$ rad and $\Delta\sigma_{\phi,\text{net}} = -0.041$ rad). While the increase in average wobble is not insignificant, the highly ordered $\beta$-sheet assemblies that are retained over time lead to a slight underestimation of the change in morphology and a large underestimation of the surface hydrophobicity increase ($\widehat{\chi^2} = 0.212, \hat{\eta} = 0.129$). The trends in average orientation ($\bar{\phi}$), orientation spread ($\sigma_\phi$), average wobble ($\bar{\Omega}$), and wobble spread ($\sigma_\Omega$) are also shown, with $\phi$ statistics exhibiting high temporal stability and Ω statistics exhibiting slightly increased dynamicity. Scalebar: 60 nm.

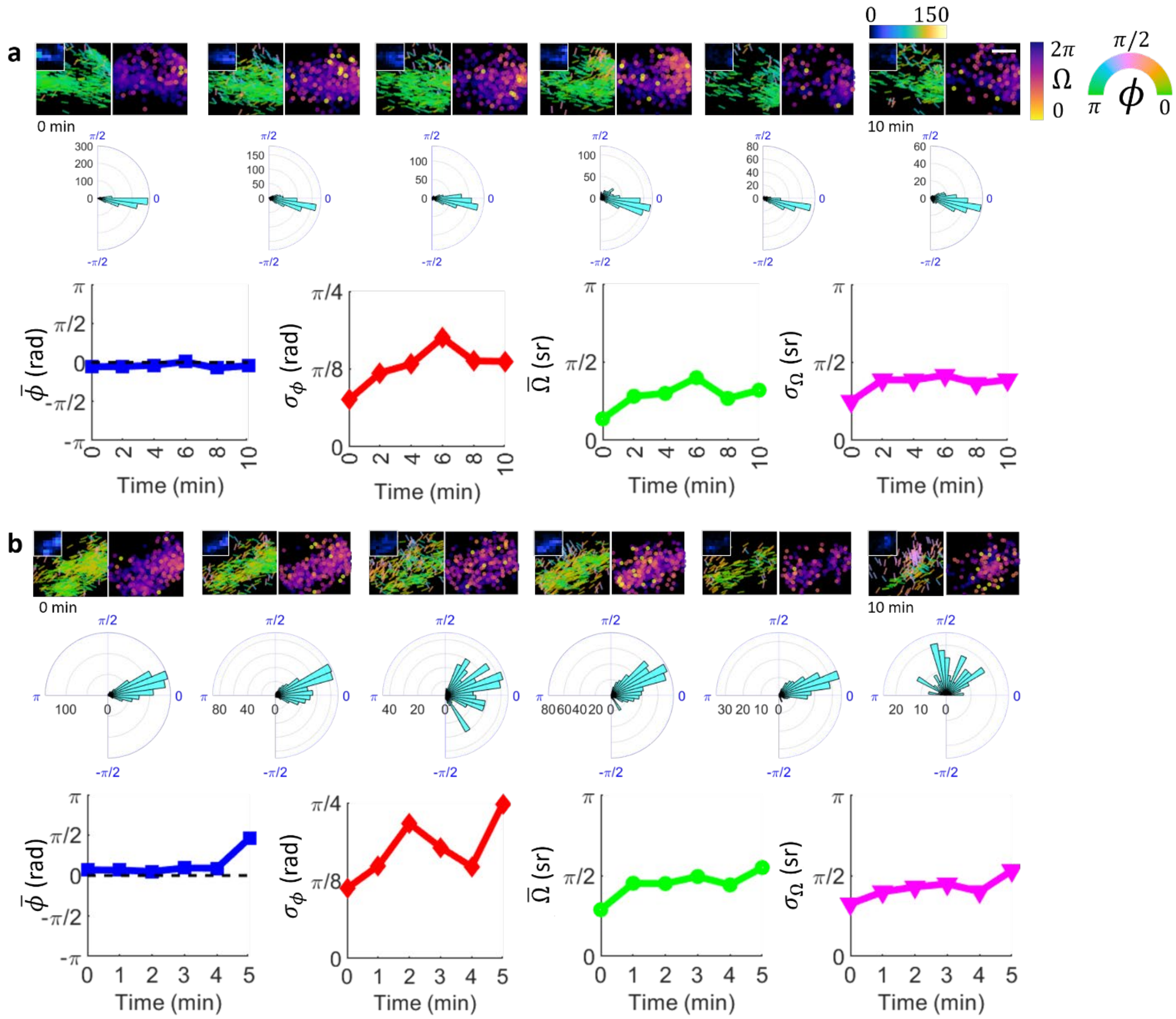

**Figure S5: SMOLM analysis of decaying Aβ42 segments**. **(a)** Uniformly decaying Aβ42 segment imaged over the course of 10 minutes. SMLM senses large morphological ($\chi^2 = 0.308$) and surface hydrophobicity ($\eta = -0.479$) changes in the segment that are, however, associated with fairly conserved $\beta$-sheet assembly organization as sensed by SMOLM ($\left|\overline{\Delta\phi_{\text{net}}}\right| = 0.042$ rad, $\Delta\sigma_{\phi,\text{net}} = +0.194$ rad). As a result, the decrease in surface hydrophobicity is noticeably underestimated ($\widehat{\chi^2} = 0.333, \hat{\eta} = -0.227$). **(b)** Haphazardly decaying Aβ42 segment imaged over the course of 5 minutes. SMLM senses moderate morphological changes ($\chi^2 = 0.266$) and greatly decreased surface hydrophobicity ($\eta = -0.586$) by way of structural disintegration. These measurements are associated with proportional increases in the degree of disorder in the β-sheet assembly ($\Delta\sigma_{\phi,\text{net}} = 0.4223$ rad) and rigidity ($\Delta\overline{\Omega}_{\text{net}} = 0.8195$ sr). As a result, the morphological and surface hydrophobicity changes are better estimated ($\widehat{\chi^2} = 0.372, \hat{\eta} = -0.447$). Scalebar: 60 nm.

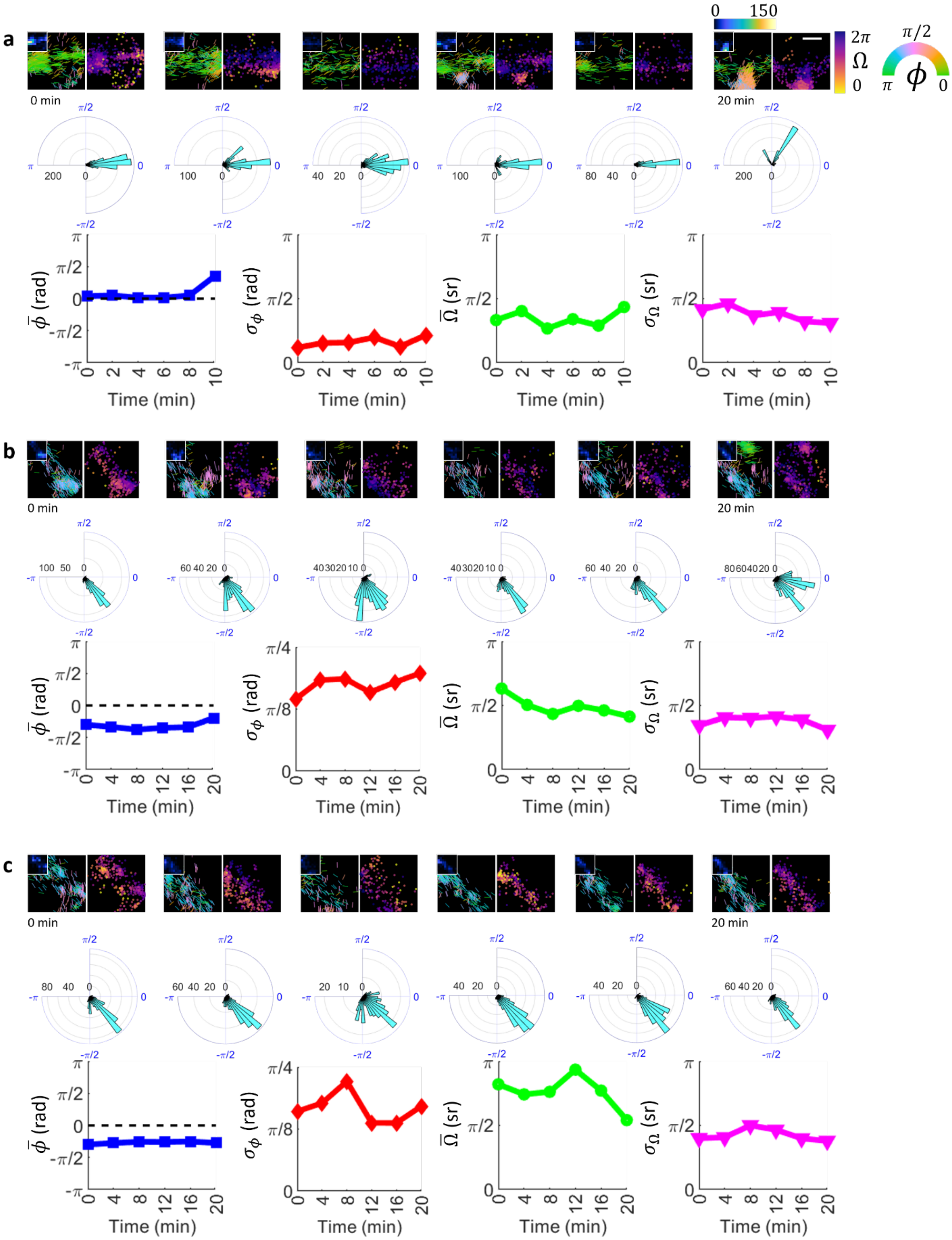
a
0 min
20 min
0
150
2π
Ω
0
π/2
π
ϕ
0
Time (min)
$\bar{\phi}$ (rad)
$\sigma_\phi$ (rad)
$\bar{\Omega}$ (sr)
$\sigma_\Omega$ (sr)
b
0 min
20 min
c
0 min
20 min

**Figure S6: SMOLM analysis of Aβ42 segments experiencing large morphological changes without significant growth and decay. (a)** Drastically remodeling Aβ42 segment imaged over the course of 10 minutes. SMLM senses large changes in shape ($\chi^2 = 0.751$) and small overall decay ($\eta = -0.128$), which are associated with large changes in underlying $\beta$-sheet orientation sensed by SMOLM ($|\Delta\phi_{\text{net}}| = 0.997$ rad). Given the large change in SMOLM statistics, the extent of morphological change is reasonably estimated, but decay is noticeably overestimated ($\widehat{\chi^2} = 0.71, \hat{\eta} = -0.304$). **(b)** Aβ42 segment with a nucleating oligomer imaged over the course of 10 minutes. SMLM senses large shape changes ($\chi^2 = 0.57$) and small overall growth ($\eta = 0.114$), which are associated with moderate changes in $\beta$-sheet orientation and small changes in $\beta$-sheet orderedness ($\Delta\phi_{\text{net}} = 0.330$ rad, $\Delta\sigma_{\phi,\text{net}} = +0.163$ rad); the nucleating oligomer features $\beta$-sheet assemblies oriented only somewhat dissimilar to the main segment. Note that both the $\beta$-sheet disorder and NB wobble are large ($\text{avg}(\sigma_\phi) = 0.550$ rad, $\text{avg}(\overline{\Omega}) = 1.540$ sr) compared to (a). As a result, the morphological change and overall growth are underestimated ($\widehat{\chi^2} = 0.405, \hat{\eta} = -0.058$). **(c)** Remodeling Aβ42 segment imaged over the course of 10 minutes. SMLM senses large shape changes ($\chi^2 = 0.539$) and small overall decay ($\eta = -0.11$), which are again associated with very small overall changes in $\beta$-sheet orientation and orderedness ($|\Delta\phi_{\text{net}}| = 0.077$ rad, $\Delta\sigma_{\phi,\text{net}} = 0.033$ rad). Note that both the $\beta$-sheet disorder and NB wobble are large ($\text{avg}(\sigma_\phi) = 0.526$ rad, $\text{avg}(\overline{\Omega}) = 2.394$ sr) compared to (a). As a result, the morphological change and overall growth are underestimated ($\widehat{\chi^2} = 0.248, \hat{\eta} = -0.32$).

### 9. SubROI SMOLM images (all timestamps) and associated statistics

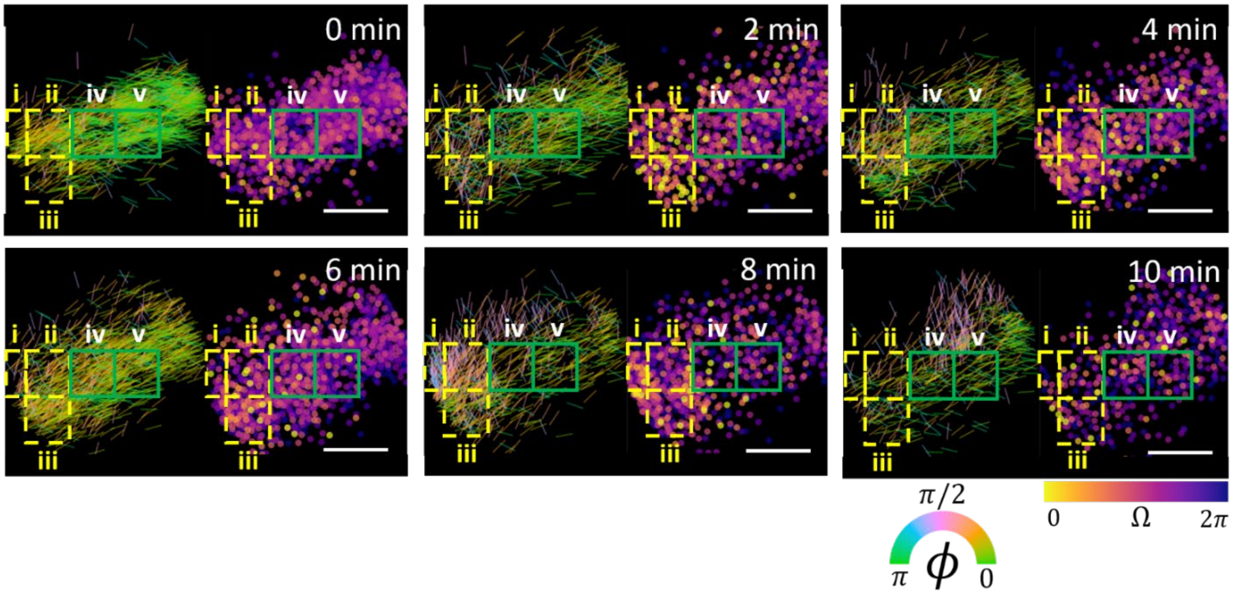

**Figure S7: SMOLM time-lapse images of remodeling Aβ42 segment shown in Figure 4a.** For each set of SMOLM images, taken at 2 min. intervals, the (left) orientation ϕ, represented as a line segment aligned parallel to its orientation, and (right) wobble Ω of each NB molecule are shown. **(i-iii)** (dashed yellow boxes) feature more disordered underlying assemblies on average than their neighboring subregions **(iv,v)** (green boxes).

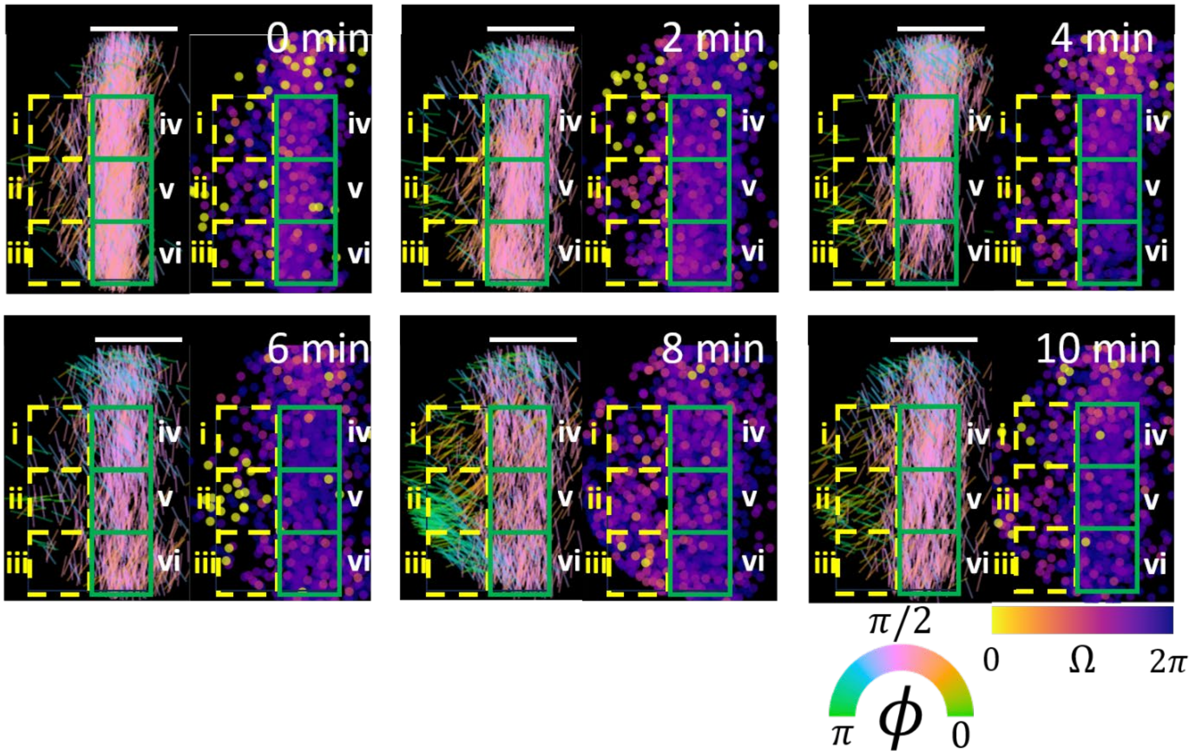

**Figure S8: SMOLM time-lapse images of remodeling Aβ42 segment shown in Figure 4b.** For each set of SMOLM images, taken at 2 min. intervals, the (left) orientation ϕ, represented as a line segment aligned parallel to its orientation, and (right) wobble Ω of each NB molecule are shown. SubROIs **(i-iii)** (dashed yellow boxes) have more disordered underlying assemblies than the core of the fibril **(iv-vi)** (boxed in green).

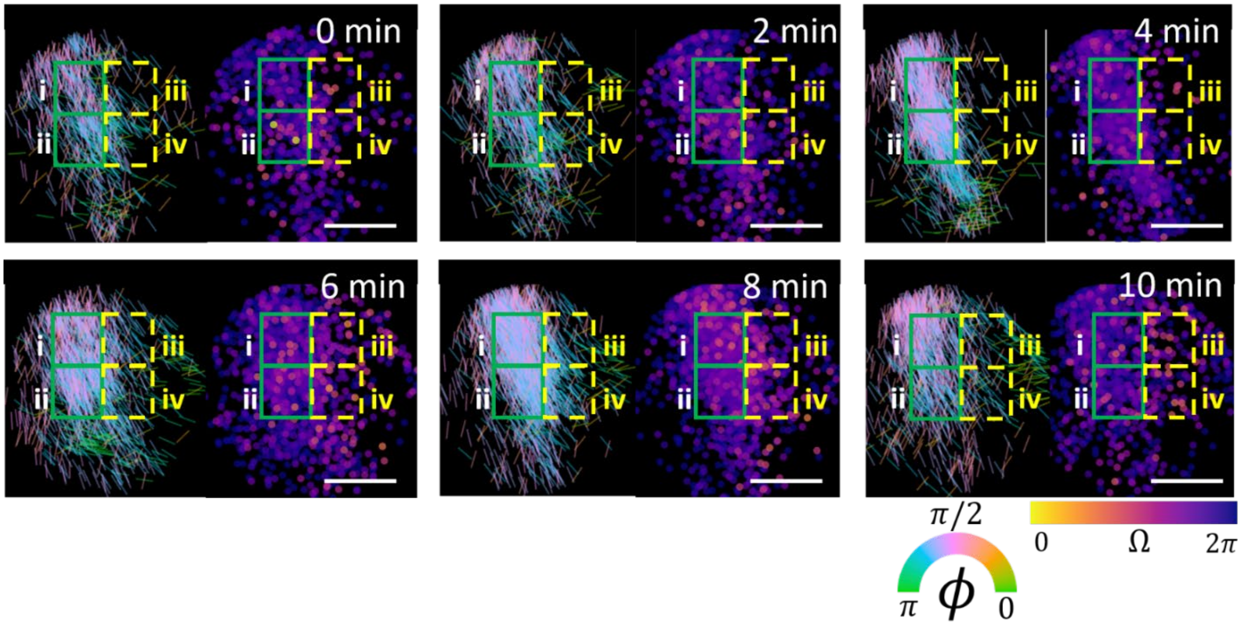

**Figure S9: SMOLM time-lapse images of remodeling Aβ42 segment shown in Figure 4c.** For each set of SMOLM images, taken at 2 min. intervals, the (left) orientation ϕ, represented as a line segment aligned parallel to its orientation, and (right) wobble Ω of each NB molecule are shown. The dynamic edges of the fibril in subROIs **(iii,iv)** (dashed yellow boxes) are more disordered and structurally heterogeneous than the segment core in subROIs **(i,ii)** (green boxes).

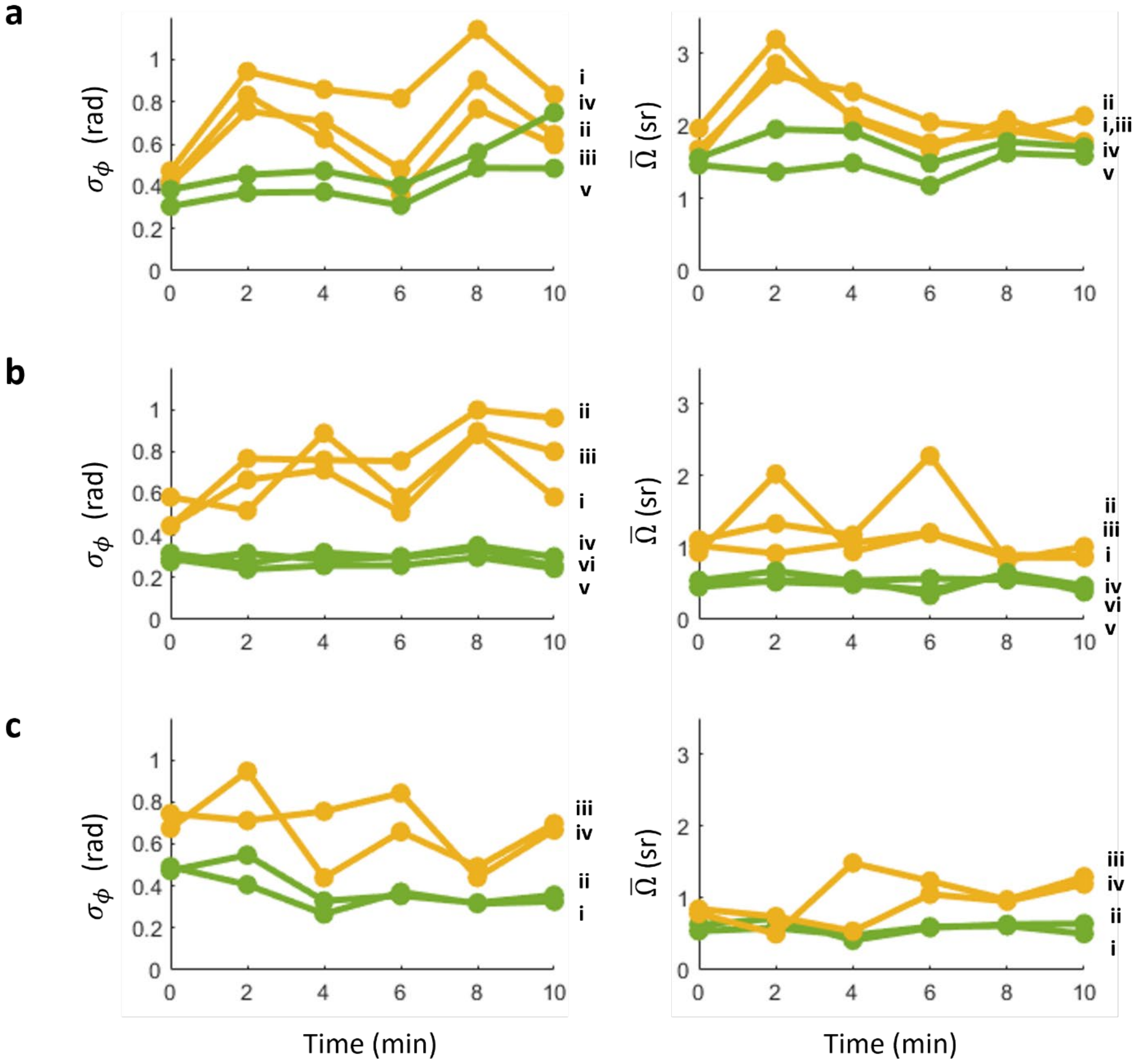

**Figure S10: Orientational uniformity $\sigma_\phi$ and NB binding rigidity $\overline{\Omega}$ over time for subROIs shown in Figures 4(a-c).** Yellow curves correspond to assemblies that are more disordered on average than their neighboring subregions (in green).

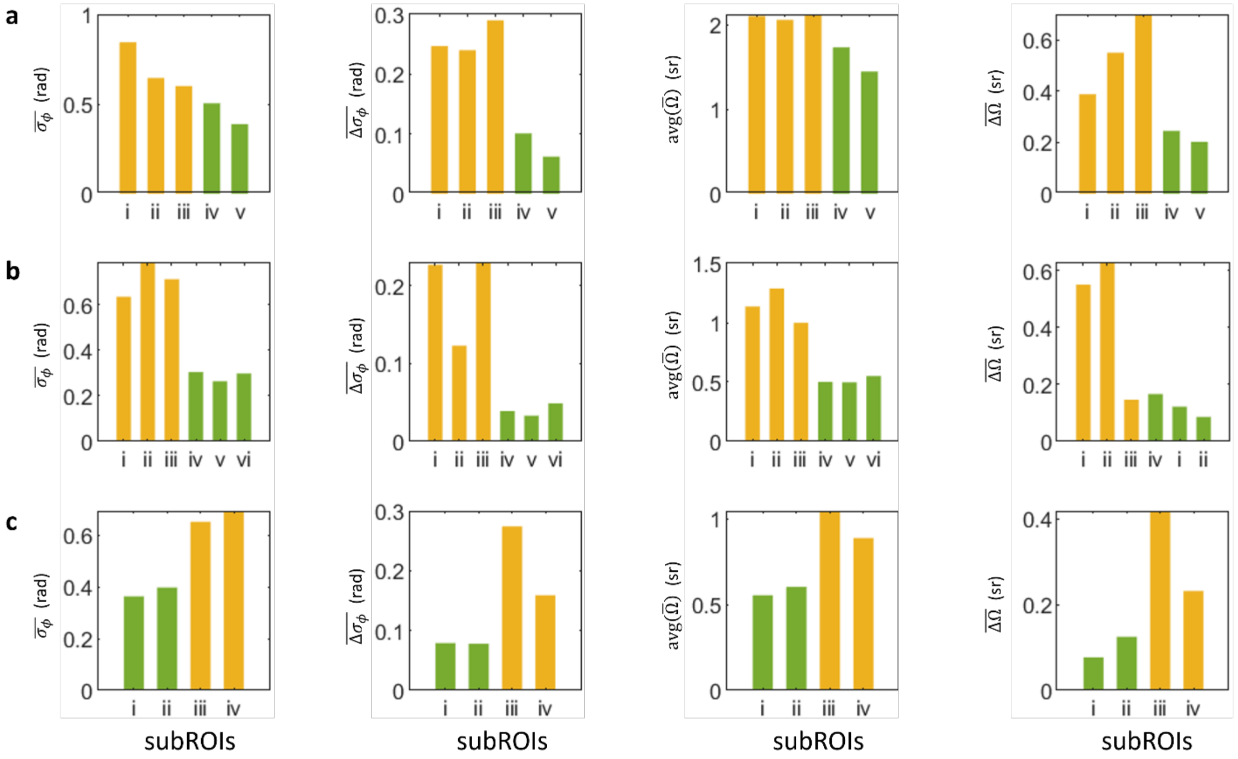

**Figure S11: Average subROI SMOLM statistics associated with Figures 4(a-c).** $\overline{\sigma_\phi}$ represents the orientation spread measured in each subROI averaged over all time points, and $\overline{\Delta\sigma_\phi}$ is as defined in Table S1. avg($\overline{\Omega}$) is the mean wobble averaged over time, and $\overline{\Delta\Omega}$ is as defined in Table S1. Yellow bars correspond to assemblies that are more disordered on average than their neighboring subregions (in green).

### 10. Supplementary Movies

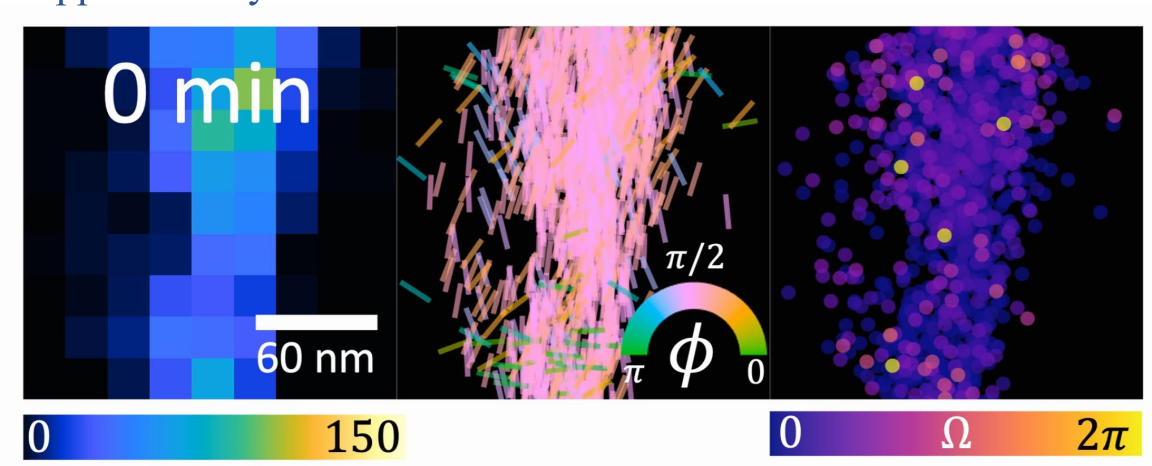

**Movie S1.** Stable Aβ42 segment measured over 10 minutes, corresponding to the ROI shown in Figure 3a. The initial SMOLM image (0 min.) visualizes the segment prior to irradiation. (Left) SMLM and (middle and right) SMOLM images of (middle) orientation φ and (right) wobble Ω. Colorbars: (Left) NB localizations per 20 nm × 20 nm bin, (middle) NB orientation $\phi$ (rad), (right) NB wobble Ω (sr). Scalebar: 60 nm.

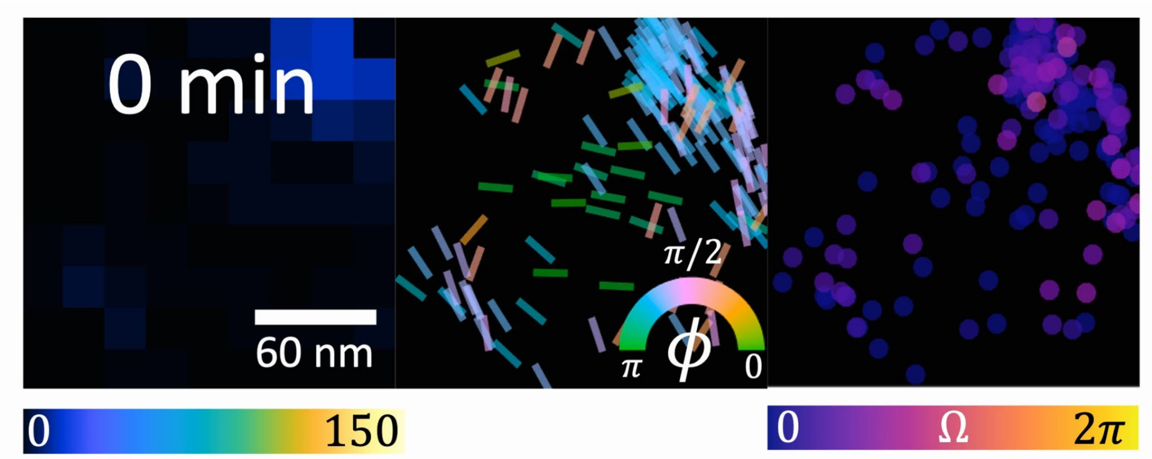

**Movie S2.** Growing Aβ42 oligomer measured over 10 minutes, corresponding to the ROI shown in Figure 3b. The initial SMOLM image (0 min.) visualizes the segment prior to irradiation. (Left) SMLM and (middle and right) SMOLM images of (middle) orientation φ and (right) wobble Ω. (Left) NB localizations per 20 nm × 20 nm bin, (middle) NB orientation $\phi$ (rad), (right) NB wobble Ω (sr). Scalebar: 60 nm.

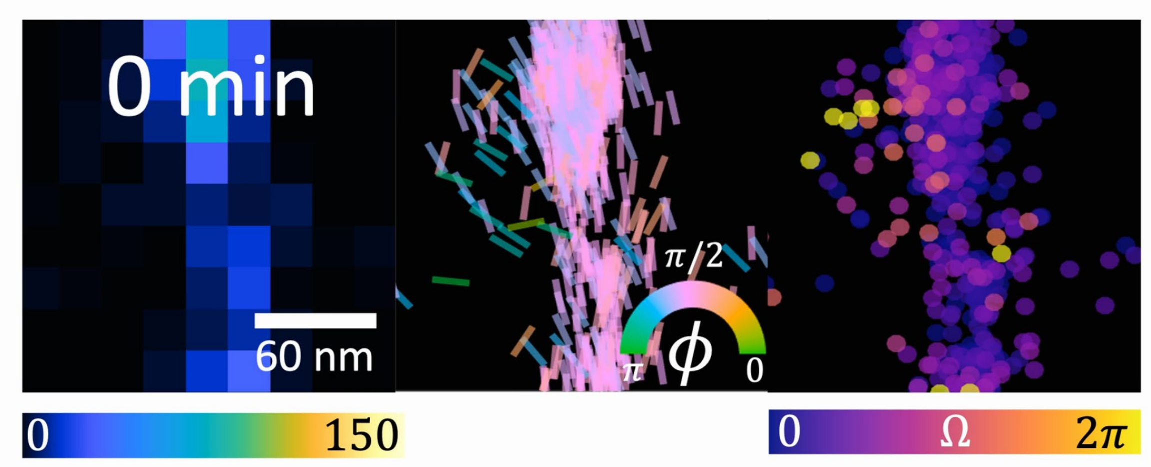

**Movie S3.** Rapidly decaying Aβ42 segment measured over 5 minutes, corresponding to the ROI shown in Figure 3c. The initial SMOLM image (0 min.) visualizes the segment prior to irradiation. (Left) SMLM and (middle and right) SMOLM images of (middle) orientation φ and (right) wobble Ω. (Left) NB localizations per 20 nm × 20 nm bin, (middle) NB orientation $\phi$ (rad), (right) NB wobble Ω (sr). Scalebar: 60 nm.
